# Germline genomic and methylomic dynamics following three generations of early-life metabolic challenges

**DOI:** 10.64898/2026.07.21.739755

**Authors:** Violeta de Anca Prado, Fábio Pértille, Dennis Andersson, Marta Mourin-Fernandez, Marta Gòdia, Josep C Jiménez-Chillarón, Joëlle Rüegg, Carlos Guerrero-Bosagna

## Abstract

Environmental and dietary factors can exert multigenerational effects on health and development. In this study, we investigated whether early-life metabolic challenge affects the germline genome and epigenome across three generations. Using a murine model of early life obesity via litter size reduction (overnutrition group, ON) and a control group (CT), we followed the paternal lineage focusing on germline genomic and methylation changes employing Genotyping-by-Sequencing (GBS) coupled with methyl-immunoprecipitation (GBS-MeDIP). We found that unrelated ON families clustered together based on identified Single-Nucleotide Polymorphism (SNP), suggesting that the treatment may have genomic impact. Copy number variations (CNVs) events were identified in ON individuals, being enriched in Long Interspersed Nuclear Elements (LINEs) and Long Terminal Repeats (LTRs). While Principal Component Analysis (PCA) of the methylome showed no clear treatment effect, pathway enrichment and regional analyses revealed methylation changes associated with transposable elements and developmental genes. Notably, the ON group exhibited a disruption in the methylation of Repetitive Elements (RE), which was significant in the same type of RE that were also enriched in the observed CNVs. The ON also showed reduced emergence of novel SNPs in offspring compared to the CT group. These findings suggest that multigenerational metabolic challenge can constrain genetic variability and induce genome instability, potentially mediated by transposable element activity rather than widespread changes in DNA methylation. This work highlights the importance of studying both genome and epigenome dynamics under realistic, multigenerational exposure scenarios and suggests that early metabolic challenges can have long-lasting impacts on genomic architecture and evolutionary potential.

## Introduction

Environmental obesogens are defined as “chemicals that initiate or exacerbate the development of obesity and their associated health consequences” (Grün & Blumberg, 2007). An obesogenic environment may include exposure to environmental toxicants, as well as dietary behaviour, inactive lifestyle and geographical units (Mei et al., 2021). Dietary behaviours may include non-usual dietary patterns in early life stages such as overfeeding (Pentinat et al., 2010) or malnutrition (Lecoutre et al., 2019) in the perinatal stage or during lactation. Consequently, mammals exposed to an obesogenic environment are at higher risk of developing obesity and metabolic disease (Grün & Blumberg, 2007), leading to increased Reactive Oxygen Species (ROS), inflammation levels (Savini et al., 2016), sperm DNA damage, as well as changes in the endocrine balance (Fariello et al., 2012); (Taha et al., 2016).

In addition to these direct effects, the development of metabolic disease during the lifetime of an individual can also affect subsequent generations. Indeed, studies on humans and mice have shown multi– and transgenerational transmission of several phenotypic markers upon metabolic challenges via the paternal line (Pentinat et al., 2010); (Jimenez-Chillaron et al., 2016). This transmission has been argued to be mediated not by genetics, but by different epigenetic mechanisms via the germline (Incedal Irgat & Bakirhan, 2022). Research shows that at least two generations can be affected by adulthood obesity via changes in the miRNA sperm load from the exposed generation (Jimenez-Chillaron et al., 2016); (Fullston et al., 2013). Another possible mechanism is via heritable epigenetic marks that overcome the early embryonic epigenetic reprogramming (Hieronimus & Ensenauer, 2021). Specifically, DNA methylation changes in sperm have been observed in mammals between individuals from unexposed descend from pregnant females ancestrally versus those exposed to a variety of insults that cause metabolic disruption, including environmental toxicants such as bisphenol A (BPA) (Manikkam et al., 2013); (Ben Maamar et al., 2023) and dichlorodiphenyltrichloroethane (DDT) (Skinner et al., 2013); (Thorson & Skinner, 2023).

These transgenerational germ line epimutations could have an impact on the genome too (Guerrero-Bosagna, 2020). One mechanism that has been discussed for this phenomena is the hypermutability of methylated cytosines, which have a twelve-fold higher probability to spontaneously deaminate to thymine compared to unmethylated cytosines (Hardwick et al., 2018). Another possible mechanism is the deregulation of repeat element activity causing copy number variations (CNV), which has been shown to be more occurrent in the paternal germ-line (Wang, 2017). The potential genomic effects derived from epigenetic changes in the germ line are not only controlled by DNA methylation but also by piRNAs (Wang, 2017). Therefore, germ-line epigenome alterations persisting across generations could induce genomic variability via SNPs and/or CNVs. Indeed, the level and populational variability of sperm dinucleotide cytosine, followed by a guanine (CpG) methylation has been shown to relate to the emergence of SNPs or CNVs in chicken breed diversification (Pértille et al., 2019).

While numerous studies have investigated the effects of obesogenic challenges across generations, they often investigate canonical transgenerational effects, for example in the developmentally exposed generation and non-exposed descendants (Nilsson et al., 2020), (Thorson et al., 2021). Although some research has been done on the effects of multigenerational exposures to different obesogenic toxicants (Nilsson et al., 2023), (Brulport et al., 2021), the long-term genomic impacts resulting from exposure affecting successive generations is poorly understood. Importantly, compared to transgenerational processes (i.e., when the effect of a single developmental exposure is transmitted across generations), multigenerational exposures represent a more realistic picture. In this realm, the dynamics between the germline epigenome and genome occurring throughout the exposed generations are very important to understand, albeit still unknown. To our knowledge, no studies have addressed how the genome is affected by sperm DNA methylation changes in a controlled experiment of multigenerational exposure. The main aim of this study was to determine if multigenerational exposure to early life metabolic challenge affects the germline methylome and genome, as well as to discern the dynamics between the methylome and the genome. We employed litter size reduction during the neonatal period in mice, as a model of early life metabolic challenge. Individuals raised in small litters developed later-in-life obesity and metabolic syndrome (Pentinat et al., 2010). This treatment was maintained across three generations in which mouse families were tracked to distinguish family from treatment effects.

## Material and methods

### Animal care and experimental design

The ICR mouse strain (ICR-CD1, Harlan Laboratories, Italy) was chosen as model due to its rapid somatic growth as described in (Pentinat et al., 2010). For three generations (F0-F2), eight-week-old naïve independent females were mated with non-sibling males. After delivery, litter size was adjusted to 8 pups (mixed sex) per dam in the control group (CT) and the treatment group consisted of 4 pups per dam in what was called the overnutrition group (ON). The pups could nurse freely and were weaned at 3 weeks old onto the standard chow. After sexual maturation at 3 months, males were mated with external naïve females. At four months old CT and ON individuals were weighted, tested their blood glucose and insulin. Upon confirmation of pregnancy for the next generation, male mice were euthanized, weighted, liver, epidydimal and inguinal white adipose tissue (eWAT and iWAT) were dissected and weighted, testes were dissected, frozen in liquid nitrogen and stored at –80 degrees for later sperm isolation. Ten physiological and metabolic parameters (BW-4M, BW-SAC, blood glucose levels, blood insulin levels, liver, eWAT and iWAT weights, and each tissue weight expressed as a percentage of body weight) were compared between the control (CT) and overnutrition (ON) groups within each generation (F1: CT n = 12, ON n = 12; F2: CT n = 13, ON n = 14). Missing or non-numeric values were excluded on a per-parameter basis. For each parameter and generation, the normality of each group was assessed with the Shapiro-Wilk test. When both groups met the normality assumption (p > 0.05), homogeneity of variances was evaluated with Levene’s test and the groups were compared using an unpaired two-tailed Student’s t-test, or Welch’s t-test when variances were unequal. When either group deviated from normality, the two-tailed Mann-Whitney U test was applied instead. Across the 20 comparisons this yielded 11 Student’s t-tests, 1 Welch’s t-test and 8 Mann–Whitney U tests. Significance was set at p < 0.05 and p-values are reported without correction for multiple comparisons, indicated as * p < 0.05, ** p < 0.01 and *** p < 0.001. All animal procedures were approved by the Committee of Animal Experimentation from the University of Barcelona and the Conselleria de Ramaderia I Pesca de la Generalitat de Catalunya, Spain. No. 10010 and 10065.

### Sperm isolation

For sperm isolation from the rest of the testis cells, slices were cut from frozen testis. Later, those slices were placed into 1 mL PBS (phosphate-buffered saline), vortexed for 30 seconds and incubated for one hour at 37 °C with constant rotation. After the incubation, samples were sonicated for 5 seconds at 60 % in ice water (705 Sonic Dismembrator, Fisher Scientific, Hampton, New Hampshire, USA) followed by vortexing for 30 seconds and centrifugation of the isolated sperm at 4000 g for 3 minutes. The pellet of sperm enriched cells was washed two times in 1mL of PBS by inversion. Subsequently, it was resuspended in 820 µL of IP buffer (prepared by mixing Na-Phosphate 100 mM (pH 7.0), NaCl 5 M, Triton X-100, and DNase free UP water).

### DNA extraction

For DNA extraction, 80 µL of DTT (di-thio-treitol, 0.1 M) was added to the purified sperm followed by an incubation of 15 minutes at 65 °C. After the incubation time, 20 µL of proteinase K (20 mg/mL) (Sigma-Aldrich, St. Louis, MO, USA) was added and a second incubation was performed for 1 hour at 55 °C in rotation. After the second incubation, 300 µL of protein precipitation solution was added (Promega, Madison, Wisconsin, USA) followed by a third incubation at 4 °C for 15 minutes. Centrifugation followed at 8000 g for 20 minutes at 4 °C, the pellet was discarded and 1mL of 100 % refrigerated isopropanol and 2 µL of glycogen (5mg/mL, Sigma-Aldrich, St. Louis, MO, USA) was added to the supernatant and inverted at least 4 times to ensure a gentle mixing. A fourth incubation was performed for 30 minutes at 4 °C followed by a centrifugation at 8000 g for 30 minutes at 4 °C. The supernatant was discarded and 500 µL of refrigerated 70% ethanol was added to the pellet. Upon mixture, a final centrifugation was performed for 10 minutes at 8000 g and 4 °C. The supernatant was discarded, and any ethanol residue was left to be evaporated. The DNA pellets were resuspended in 150 µL of distilled water once the Eppendorfs were dry.

### GBS and GBS-MeDIP library

DNA integrity was verified by agarose gel electrophoresis. DNA purity was assessed using NanoDrop 2000c spectrophotometer (Thermo Fisher Scientific, Waltham, MA, USA), and DNA concentration was quantified using Qubit dsDNA High Sensitivity (HS) (Thermo Fisher Scientific, Waltham, MA, USA). After DNA quality assessment, Genotyping-By-Sequencing (GBS) and Genotyping-By-Sequencing coupled with Methyl-Immunoprecipitation (GBS-MeDIP) libraries were done as previously described (Rezaei et al., 2022). Later, quality control of both types of libraries was done using Tapestation High Sensitivity D1000 kit (Agilent Technologies). After the final quality control, 45 libraries were sent out to be quantified, clustered, and paired end sequenced in the Illumina HiSeq2500 platform with a read length of 125 bp at the SNP&SEQ facilities of the SciLifeLab (Uppsala, SE).

### Genetic analysis

#### Preprocessing of GBS

Demultiplexing of the library was performed with Stacks v.2.62 (Catchen et al., 2013), trimming of adapters and filter by base quality above 30 was done with cutadapt v.4.0 (Martin, 2011).

After the removal of low quality reads, alignment was performed on the filtered reads with bwa-mem v.2.2.1-20211213-edc703f (Li & Durbin, 2009) against the reference genome mm39 of *Mus musculus* (Clawson et al., 2023). Filtering of the reads to only select unique mapped reads was performed by filtering out reads with the tags XA and SA in the bam files using samtools v.1.14 (Danecek et al., 2021) and tags were set for downstream analysis: @RG\tID:GBSICR\tSM:<sample_ID>\tPL:ILLUMINA.

#### SNP calling and filtering

The first step for SNP calling following GATK v.4.2.0.0 best practices (Poplin et al., 2017) is to perform a recalibration step. Recalibration was performed with the SNP database from Ensembl (Harrison et al., 2024) using BaseRecalibrator (Poplin et al., 2017) and ApplyBQSR (Poplin et al., 2017) with depth (DP) > 101 and maximum DP of 1644 (corresponding to the median + standard deviation).

After recalibration, variant calling was performed per individual using HaplotypeCaller (Poplin et al., 2017), the filters GATK advised were applied (FS > 60, QD < 2.0, MQ < 40.0) with custom filters including: depth (DP) > 20, maximum depth of 1644, allele depth (AD) > 10, and a maximum of 3 called SNP every 50 bp.

After the filtering, each individual was merged into one VCF using CombineGVCFs (Poplin et al., 2017) and the genotypes were called with GenotypeGVCFs (Poplin et al., 2017). SNPs that were not present in every sample were deleted.

#### Principal Component Analysis on GBS data

To check any association between depth of sequencing and number of SNPs identified, and to evaluate the impact of the treatment onto the germ-line genome a Principal Component Analysis (PCA) was performed with plink v.1.90b4.9 (Chang et al., 2015), visualization done with ggplot2 v.3.5.1 (Wickham & Wickham, 2016) in R v.4.3.1.

To evaluate the impact of the family and the treatment in the genome the merged VCF file containing all the individuals was imported to R with the package vcfR v.1.15.0 (Knaus & Grünwald, 2017). With the resulting genotype matrix, a PCA was performed with FactoMineR v.2.11 (Lê et al., 2008). To FactoMineR the SNPs were supplied as the active variables in the PCA, and the variables family and treatment were supplied as supplementary variable. To evaluate the impact of the family and treatment, a correlation was checked between both variables with the principal components selected, using Kruskal-Wallis for statistical significance, as the coordinates did not follow normality (McKight & Najab, 2010).

#### Copy number variation, false positive control and variant prediction

Copy number variation analysis was performed using GATK v.4.3.0.0 for germline copy number variants (gCNVs) discovery (Poplin et al., 2017). The genome was binned in 1000 bp intervals with PreprocessIntervals. Afterwards the reads of each individual were assigned to each bin with CollectReadCounts. Filtering of bins with extreme counts was performed using FilterIntervals set to default mode. The next step was ploidy calling of chromosomes with DetermineGermlineContigPloidy using the CT samples, with the prior probabilities set as 0.01, 0.01, 0.097, and 0.01 for the autosomes, 0.01, 0.49, 0.49, and 0.01 for the X chromosome and 0.5, 0.5, 0.0, and 0.0 for the Y chromosome. The output created a model. Ploidy calling of ON samples were run with DetermineGermlineContigPloidy specifying the model with the flag –-model.

CNV calling was performed on the CT samples using GermlineCNVCaller using the flags –– intervals for restricting the call to the filtered intervals, –-contig-ploidy-calls for input of the CT ploidy calls, and –-run-mode set to COHORT. The output was a model for CNV calling. Afterwards ON samples were CNV called using GermlineCNVCaller specifying the flags –-model for the output model, –-contig-ploidy-calls for the ploidy call, and –-run-mode set to CASE. Finally, CNV calls of the ON samples were segmented call copy number states using PostprocessGermlineCNVCalls, specifying the flags –-calls-shard-path for the CNV call, –– contig-ploidy-calls for the ploidy call, –-model-shard-path for the CNV model, and –-allosomal-contig chrX and chrY for the sex chromosomes. The output is a vcf file with the locations of the CNV and their state, DUP for duplication or DEL for deletion.

To control for false positives in the CNV calling and due to limitations of the GBS method to detect CNVs (Lemay et al., 2019), there was a CNV calling using randomized data. In essence the CT.bam files were merged and the percentage of reads from each CT.bam that added to the total was calculated. A random subsampling using the mentioned percentage of reads was performed to simulate an equivalent situation using the command samtools view –s from samtools v.1.20 (Danecek et al., 2021). Using this randomized data all models were calculated and CNV were called on the CT samples. Those CNV are false CNVs, their characterization was useful to filter out false CNV calls in the ON run.

Variant effect prediction was performed using the program Variant Effect Prediction (VEP) (McLaren et al., 2016) using –-assembly GRCm39, –-species mus_musculus, and –-format ensembl. Annotation of repeated elements on the CNVs was performed with bedtools intersect (Quinlan, 2014). Afterwards a permutation test was performed to test for enrichment or depletion of RE in the identified CNVs. The permutation was performed with 1000 fragments of equivalent length as the CNV detected, to create the random distribution with bedtools shuffle (Quinlan, 2014). Afterwards, the fragments were annotated in the same way and empirical p-values were calculated. Visualization was done in R v.4.3.1.

#### Site Frequency Spectrum

Site Frequency Spectrum was performed to observe the proportion of shared alleles in each position among the individuals. In order to perform the Site Frequency Spectrum (SFS) test, the tool angsd v.0.933 (Korneliussen et al., 2014) was used for genotype calling, which was performed on the aligned data with flag –GL 2. After the allele frequencies were calculated with realSFS v.0.933 (Korneliussen et al., 2014) with a maximum of iterations of 1000. Visualization was done in R v.4.3.1.

#### Hierarchical tree

A hierarchical tree was built for observing genetic similarities between individuals without assuming family structure. In order to build a hierarchical tree, the genotype matrix had to be imported to R using vcfR v.1.15.0 (Knaus & Grünwald, 2017). Using the genotype matrix we calculated the Identity by State (IBS) on all our individuals using the R package SNPRelate v.1.34.1 (Zheng et al., 2012). IBS is used to determine similarity between two unrelated individuals. Later it was transformed into a dissimilarity matrix by subtracting the IBS matrix to 1. To be able to perform hierarchical clustering we used the R package fastcluster v.1.2.6 (Müllner, 2013). Hierarchical tree was plotted with R base package.

### Epigenomic analysis

#### Preprocessing of GBS-MeDIP

Demultiplexing was performed using Stacks v.2.62 (Catchen et al., 2013), after a trimming step was done for adapters and polyGs using cutadapt v.4.0 (Martin, 2011). Alignment was performed with bowtie2 v.2.3.5.1 (Langmead et al., 2019) against the reference genome mm39 of *Mus musculus* (Clawson et al., 2023), followed by filtering with samtools v.1.14 (Danecek et al., 2021) MQ > 10 in the bam files.

In order to perform differential analysis, a count matrix was created. The locations in the genome with enrichment of methylated signals were observed with MACS v.3.0.0b1 (Zhang et al., 2008). The program was ran using the flags –t for GBS-MeDIP, –c for inputting as control the GBS, –f set as BAMPE, and using the –B flag. Later, counts on the windows were calculated with featureCounts from the package subread v.2.0.3 (Liao et al., 2014), with the flags –p for paired end, –B for properly aligned and –D for counting only fragment length < 1500 bp. All the individuals were later merged using bedtools v.2.29.2 (Quinlan, 2014) into a count matrix to perform statistical tests on them.

#### Differential methylated regions (DMRs)

For performing differential methylation analysis, the count matrix was filtered so every window had a minimum of 2 individuals R v.4.3.1. The analysis was done comparing intragenerationally (CT versus ON) and intergenerationally (F1 versus F2). In the intragenerational analysis, 301 windows were analysed in F1 and 373 windows for F2 generation. For the intergenerational analysis 375 windows were analysed in CT and 315 windows in ON. Mann-Whitney was chosen as the statistical test from a recent benchmarking (de Anca Prado et al., 2026).

#### Principal Component Analysis on GBS-MeDIP data

In order to evaluate the effects of the metabolic challenge on the methylome, PCA was performed using normalized methylated windows with the package FactoMineR v.2.11 (Lê et al., 2008). Two individuals were excluded (F1.6 and F2.19) as they were outliers. The counts were supplied as active variables, and family belonging and treatment group were supplied as supplementary variables. Visualization was done in R with factoextra v.1.0.7 (Kassambara, 2016). To observe the impact of the family and the treatment in the methylome, a correlation was performed between each one and the principal components selected, using Kruskal-Wallis for statistical significance, as the coordinates did not follow normality (McKight & Najab, 2010).

#### Difference in methylation patterns

Density plots were done using the normalized methylated window counts with the package ggplot2 v.3.5.1. Annotation of the methylated windows against repeated elements was performed with bedtools intersect. Visualization was done in R v.4.3.1.

#### Pathway analysis

For pathway enrichment analysis, all the annotated regions from each individual were merged by groups and by generations. Using PathBIX (Castresana-Aguirre et al., 2021) and KEGG (Kanehisa et al., 2007) database, pathway enrichment analysis was performed with a network cutoff = 0.8 and clusterization. Visualization was done with ggplot2 v.3.5.1 (Wickham & Wickham, 2016) in R v.4.3.1.

### Annotation of GBS-MeDIP and GBS data

All the concordant pairs of reads from the GBS and GBS-MeDIP libraries were annotated using bedtools v.2.29.2. The reads were classified into different groups based on their genomic location relative to the annotation from mm39. Concordant pairs of reads within 2kb of the transcription start site (TSS) of an annotated gene were classified as “TSS + promotor region”. Concordant pairs of reads overlapping a coding sequence (CDS) were classified as “exon”. Concordant pairs of reads overlapping the 5’ untranslated region (UTR) and the start codon were classified as “5’ UTR”. Concordant pairs of reads overlapping the 3’ untranslated region (UTR) and the stop codon were classified as “3’ UTR”. Concordant pairs of reads overlapping none of the regions above described (exon + 5’ UTR + 3’ UTR + TSS + promotor region) but within a gene body were classified as “intron”. Concordant pairs of reads which did not overlap with any gene body nor a TSS+ promotor region were classified as “intergenic”. Concordant pairs of reads overlapping repeated elements (RE) were classified by the type of RE they overlap with. Concordant pairs of reads overlapping any RE were classified as “non-repeated regions”.

### Statistical test on annotated RE

After annotation of RE in the GBS-MeDIP libraries, Shapiro-Wilk tests were performed for assessing if the methylated counts of the RE followed a normal distribution. As a marginal subset only followed normality, the statistical test used to assess if there were differences on the levels of methylation between ON and CT per generation was Mann-Whitney.

### Genetic and epigenetic dynamics

Families from CT and ON across 2 generations were tracked to evaluate if any correlations could be drawn from the methylation level and the SNP emergence in the next generation. From the count matrix with the normalized counts, the genomic coordinates of all the homozygous cytosine were extracted using snpEff v.5.2 (Cingolani, 2022) in fathers. Afterwards those same locations were subset if there was an alternative allele in the GBS library of the sons using snpEff. A correlation using Spearman rank correlation test between the levels of methylation in fathers and the number of novel SNPs in offspring (homozygous SNP not inherited from the father) was done in R v.4.3.1.

For the CNV events in offspring and methylation levels in fathers, Spearman rank correlation test was performed between the overlap of methylated windows in fathers with the CNV events in their offspring in R v.4.3.1.

## Results

In this study, we investigated the sperm genomic and methylomic effects of a three generation long exposure to a metabolic challenge by litter size reduction (overnutrition group, ON; 4 pups per litter), contrasting them with a control group (CT; 8 pups per litter). For this, we employed the GBS-MeDIP method, which interrogates the genome (GBS libraries) and the methylome (GBS-MeDIP libraries) in parallel in a fraction of the genome. Analyses were conducted to determine if the sperm genome responded to the multigenerational challenge in terms of SNP and CNV changes, and whether there was a correlation between DNA methylation changes and SNPs and CNVs. We also employed novel ways to investigate sperm DNA methylation changes across generations.

### Metabolic challenge effects

A metabolic challenge was performed for three generations on ON individuals by litter size reduction. The murine model is described to trigger neonatal obesity, which is maintained throughout the rest of the adult life of the individual and which will later in life trigger metabolic syndrome related issues, such as higher insulin level in blood and higher adiposity in ratio with the body weight. All males from ON had higher body mass at the moment of sacrifice, in F2 males from the ON has significantly more iWAT compared to the CT group and in F1 ON males had a bigger liver than CT males (Figure 1). We do not observe incidence on insulin or glucose as these metabolic changes tend to unveil later in life.

**Figure 1.**
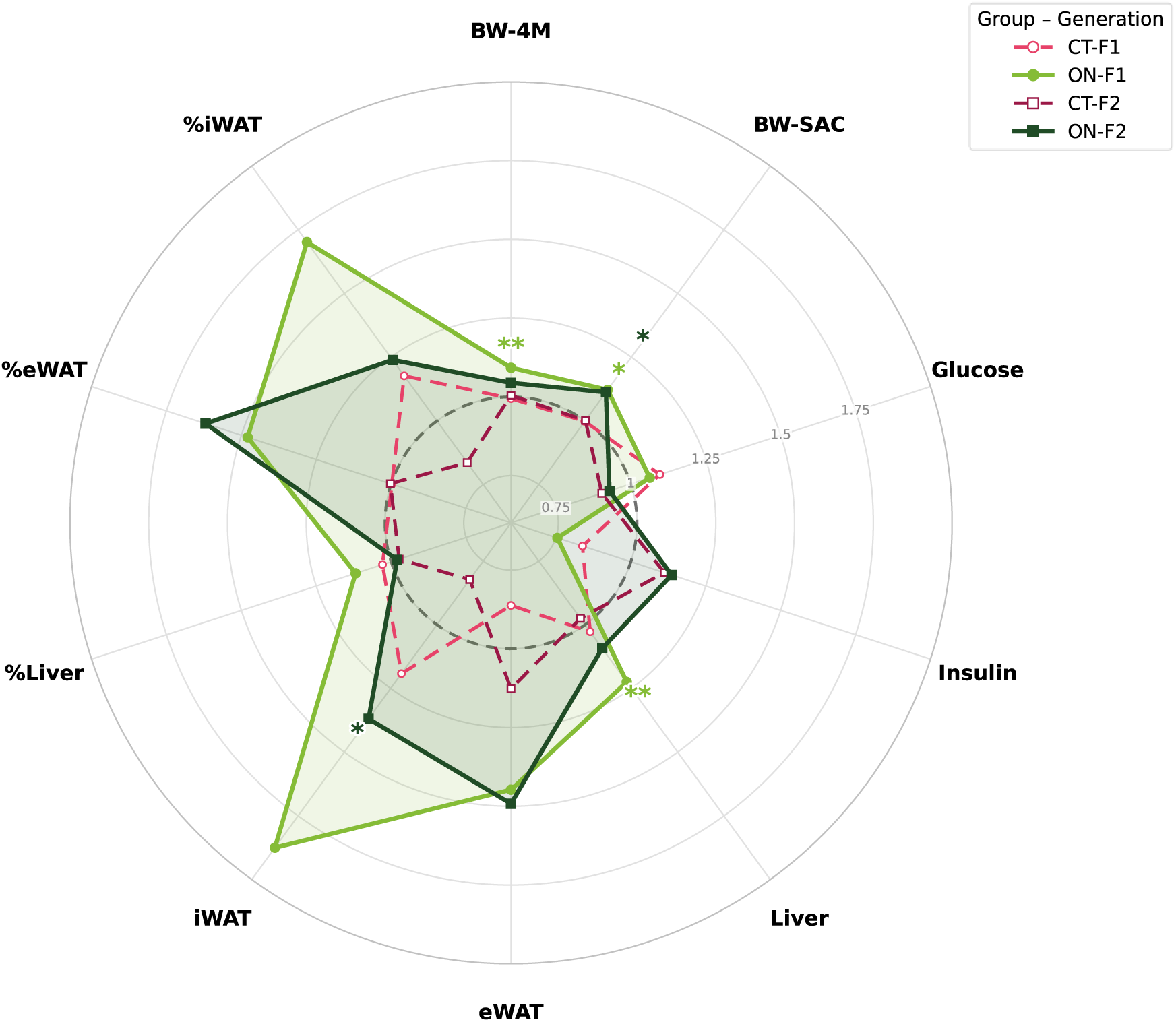
Radar plot summarizing ten physiological and metabolic parameters in male ICR mice from the control (CT) and overnutrition (ON) groups in the F1 and F2 generations (CT-F1, n = 12; ON-F1, n = 12; CT-F2, n = 13; ON-F2, n = 14). Each axis shows the group mean expressed as fold-change relative to the pooled control mean for that parameter, so the dashed ring at 1.0 represents the control reference. Parameters: BW-4M, body weight at 4 months of age measured in fed animals; BW-SAC, body weight at sacrifice measured after an overnight fast; Glucose, blood glucose (mg/dL); Insulin, blood insulin (mU/L); Liver, liver weight (gr); eWAT, epididymal white adipose tissue weight (gr); iWAT, inguinal white adipose tissue weight (gr); %Liver, liver weight relative to body weight; %eWAT, eWAT weight relative to body weight; %iWAT, iWAT weight relative to body weight (tissue weights in grams). CT versus ON differences within each generation were tested per parameter after assessing normality with the Shapiro-Wilk test, using an unpaired t-test when both groups were normally dis-tributed, or the Mann-Whitney U test otherwise; p-values were not corrected for multiple comparisons. Asterisks mark significant CT vs ON differences. Significant differences were detected for BW-4M, BW-SAC and liver weight in F1, and for BW-SAC and iWAT weight in F2.

### Basic sequencing parameters

Sequenced GBS libraries (N = 45) generated 317M reads, averaging 5,664,761 reads per individual. Mapping on average 0.3% of the mouse genome per individual, an expected coverage from GBS. A total of 2,203,415 SNPs were identified (FS > 60, QD < 2.0, MQ < 40.0, 20 > DP < 1644, AD > 10, maximum of 3 called SNPs every 50 bp) among 45 individuals, with an average of 39,347 per individual. Sequenced GBS-MeDIP libraries generated 33M reads, with an average of 589,834 reads per individual, mapping on average 0.03% of the mouse genome per individual. A total of 13,426 windows was identified across the 45 GBS-MeDIP libraries, using the GBS data from the same individual as the genomic background (min 100 bp, max 1,500 bp). Individual statistics of the metrics above are shown in Supplementary Table 1.

### Genetic results

#### Founders effect robustness of the murine model

In a population with founder effect, genetic diversity is expected to be low with high percentage of shared alleles among individuals. Consequently, results derived from studying such populations may lack relevance. Therefore, we investigated confounding factors that could emerge from the starting size of 10 individuals in F0 in both CT and ON. We employed the Site Frequency Spectrum (SFS) test, which measures the proportion of shared alleles in each position among the population, categorized by group in each generation. A population with high number of shared alleles signals founder effects, as populations that share a high percentage of SNPs do not have a high genetic variability. In our population, we observed a high frequency of singletons (SNPs which are not shared among individuals), meaning the populations is in expansion and possesses genetic diversity, with a high frequency of alleles not shared among individuals (Supplementary Figure 1). The reason for this high genetic diversity is likely due to the fact that the ICR strain is outbred, and genetic novelty was introduced by new mothers in each generation and in each group. Thus, our analysis showed that the genetic diversity of the two groups investigated (CT and ON) was not a cofounding factor in the results presented in this paper.

#### Impacts of metabolic challenge across three generations on the germline genome

From the GBS libraries, we identified a set of 33,906 SNPs against the Ensembl SNP database of *Mus musculus*, having on average 5,558 SNPs in ON and 5,516 in CT per individual. To determine if the metabolic challenge affected the sperm genome, we first performed a PCA on the identified 33,906 SNPs, as shows genetic similarity between individuals. This analysis showed clustering of all the CT families, and two distinct clusters formed by different ON families (Figure 2). One of these clusters was composed of one single family, while the other (upper left cluster in Figure 1) consisted of two families. Thus, the latter cluster was showing genetic similarity of two unrelated families, suggesting that the treatment posed an impact (otherwise the families would cluster independently, as the members of each family are genetically closest). To exclude that sequencing coverage drives the PCA clustering, we also conducted the analyses taking sequencing depth into account (Supplementary Figure 2). Similar clustering among all groups was observed, showing that sequencing depth was not a driving factor in the PCA.

**Figure 2.**
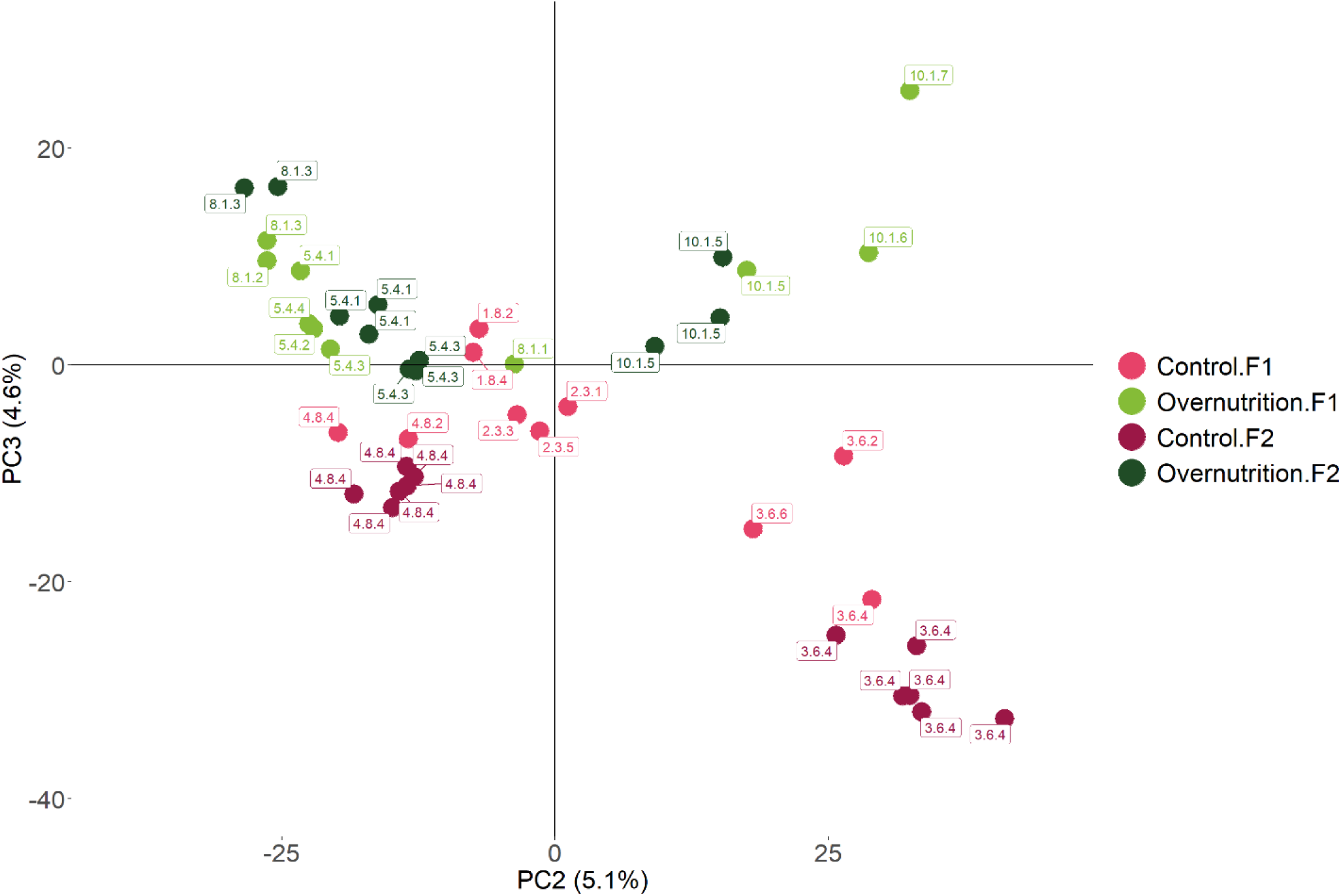
Principal Component Analysis on genetic data derived from GBS. In shades of pink, we find the two different generations of the control group (CT). In shades of green we find the two generations belonging to the overnutrition group (ON). Each individual is marked with their family name, in total there are seven families (ON = 3, CT = 4). There is a clustering of two unrelated families in ON, 8.1 and 5.4, supporting the hypothesis that the metabolic challenge posed an effect on the genome of ON individuals. On the contrary, CT families do not form clusters of individuals beyond the family relatedness.

To explain the clustering observed of unrelated families (Figure 2), we assessed the impact of treatment and family belonging with two different approaches, by performing an association test between each of the variables and each PC and by the formation of a hierarchical tree using IBS. A strong association between either family or treatment with the PC indicates that particular variable explains the variability of that PC. Table 1 presents the R^2^ and the p-values associated with each principal component (PC) and the treatment/family. The treatment is slightly correlated with the third principal component, whereas the family is highly correlated with both PCs. This slight association indicates that the treatment partially explains 4.6% of the genetic variance.

**Table 1.**
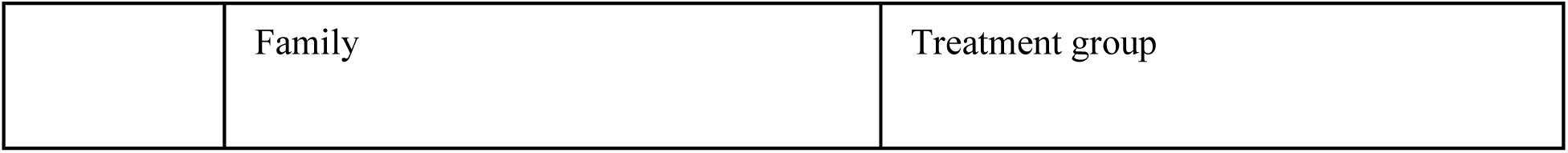
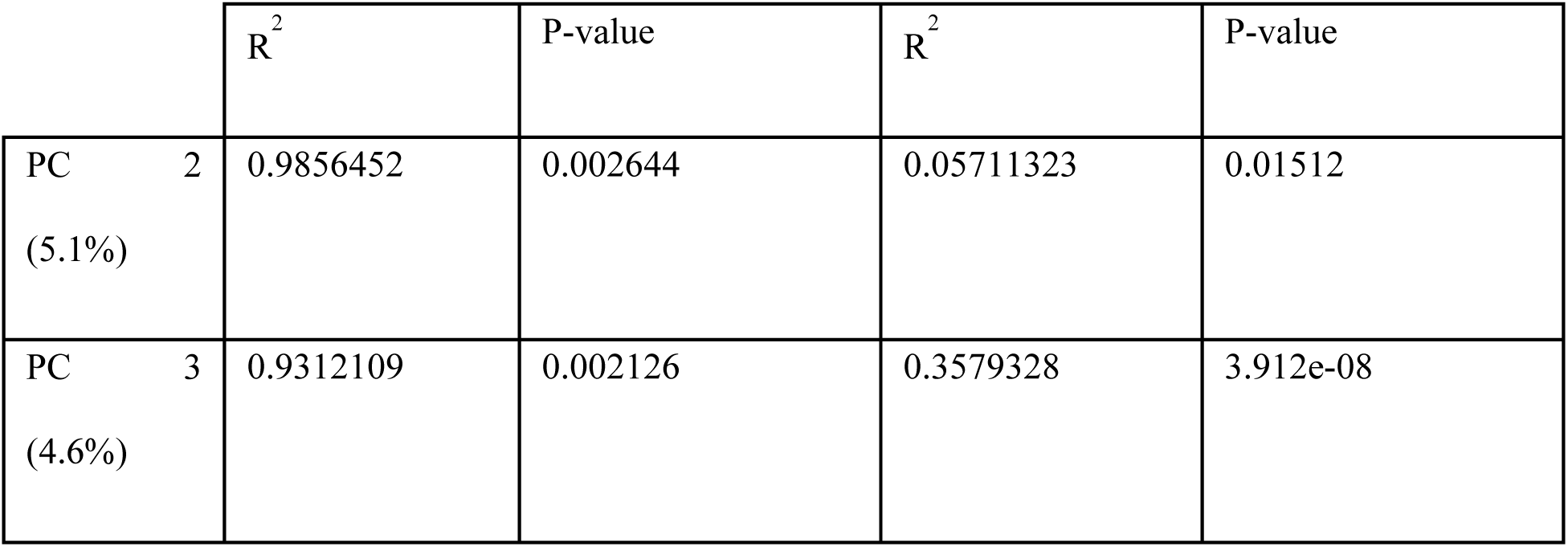
Association of treatment and families with the principal components of the PCA of the genomic data. As the data did not follow normality, Kruskal-Wallis test was used to assess statistical significance of the correlations.

The construction of a hierarchical tree using IBS assumes no family relatedness, with this approach, if the treatment had not posed any impact, only the individuals belonging to each family would create branches, and each of the families would be branching separately, as we found high genetic diversity in our initial analyses. The tree showed the formation of two main branches, each of them containing secondary branches with clear separation of CT and ON individuals, with a particular branch created from unrelated ON families (Figure 3). This reinforces the hypothesis that the treatment impacts the genome, as non-related individuals are clustered together showing a higher genetic similarity than when compared with individuals of further branches.

**Figure 3.**
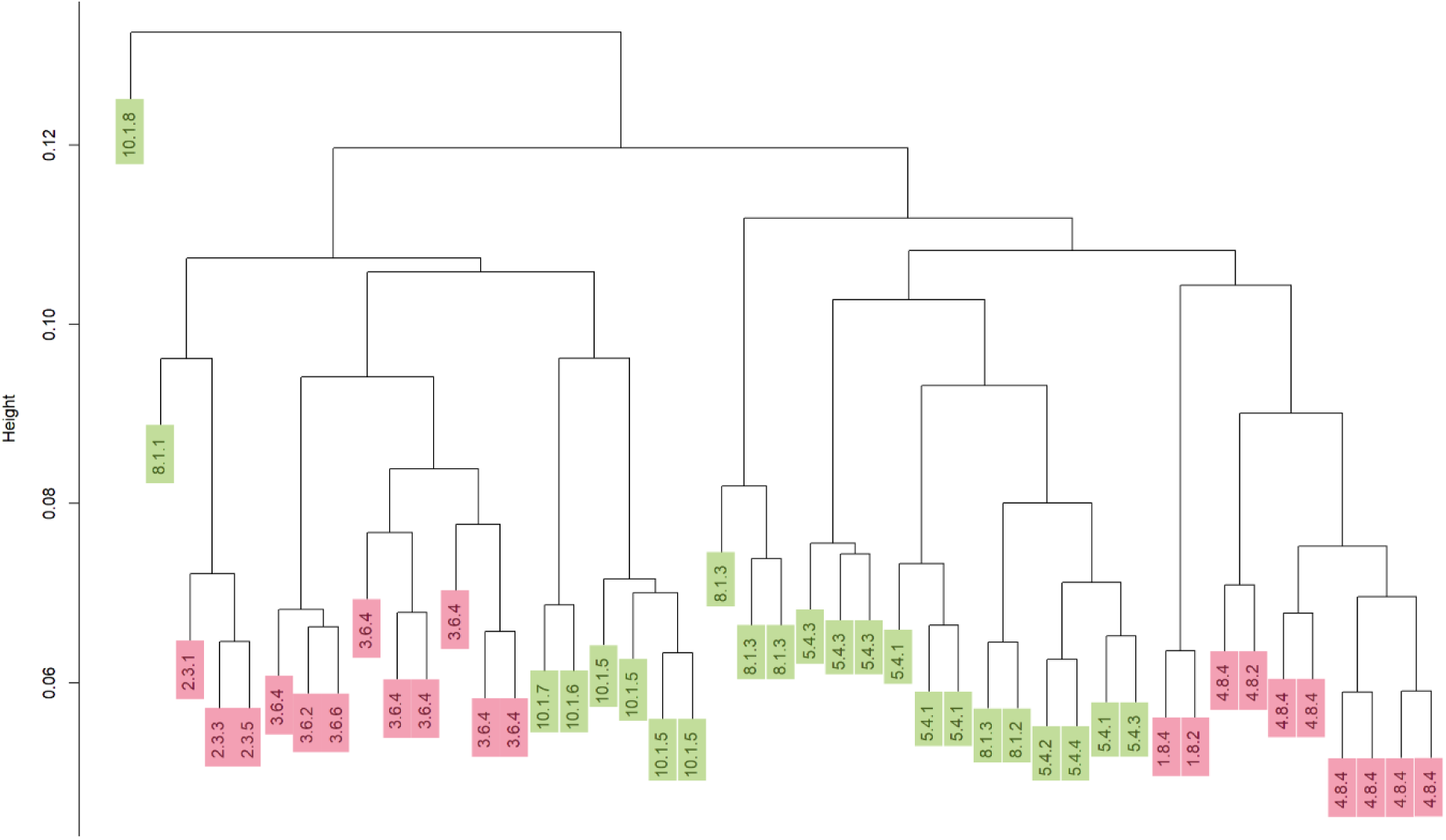
Hierarchical clustering from Identity by state (IBS) matrix derived from SNP data. All the individuals (F1 and F2) are named by their family code, ON individuals are depicted in green, CT in pink. The formation of two main branches is observed, each containing subbranches with single families in the case of CT individuals. When observing the formation of branches in ON families, is observed the formation of a branch composed of individuals from the 5.4 and 8.1 families, suggesting that, with the assumption of no blood relationships, these unrelated individuals are closer genetically.

#### Treatment effects on the emergence of CNVs

Since obesity has been linked to genome instability, we also evaluated if putative CNVs events were produced by the early life metabolic challenge. For this, first a quality control step was performed using a random model created by random subsampling of all CT reads. This analysis confirmed that most false positives CNV events are shared by fewer individuals (< 3/26) and were large events (> 1 Gbp). Based on the literature and the quality check on our own data, we filtered by: (a) minimum size (> 50,000 bp) (Lemay et al., 2019), (b) amount of individuals sharing an event (> 50%), and (c) maximum size of the CNV event (< 1Gbp). Afterwards CNV were defined in the ON individuals in comparison to the relative abundance of reads in CT individuals in a region. This resulted in the identification of 7 putative CNV events in F1 ON that were maintained in F2, 3 of which mapping to chromosome 4 (Figure 4). The 4 deletions identified over generations are maintained with the same boundaries, whereas the 3 duplications events are either expanded over generations (in the case of the duplications in chr4 and 8) or the boundaries are not maintained over the generations.

**Figure 4.**
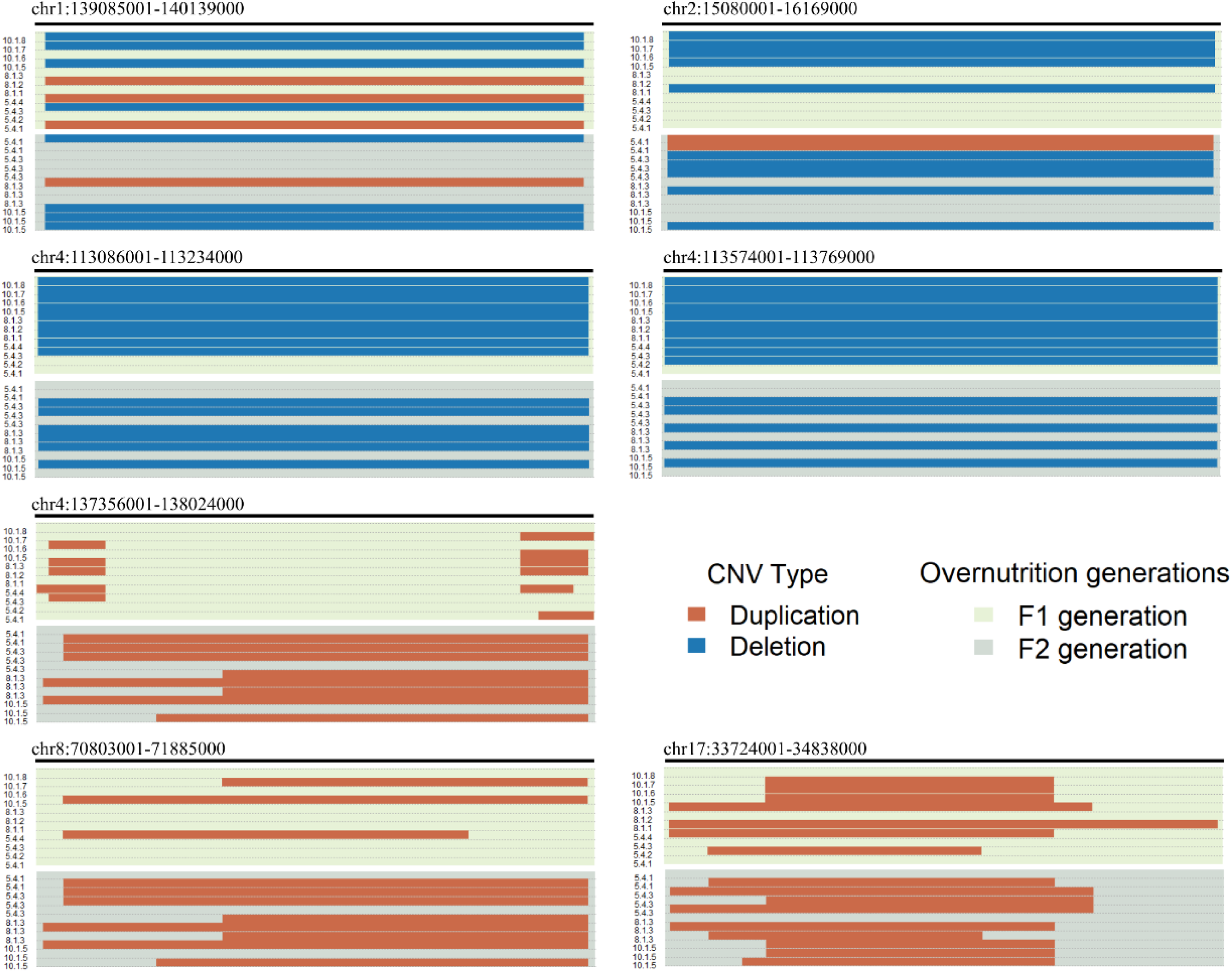
Position and length of the CNV events per individual observed in ON in the generations F1 and F2. Individuals are only marked in the Y axis with the family code. Duplications are shown in orange whereas deletions are shown in blue. Chromosome 4 is the most affected, with three CNV events expanding more than 20.000bps per CNV. While deletion is stable over generations, duplication events seem to be expanded in F2, less stable.

Those putative CNV events were annotated using VEP in order to determine their potential consequences. In the F1 generation, 46.2% of the CNV were predicted to lead to transcript amplification, 26.9% to feature truncations, 19.2% to loss of transcriptional stop and 7.7% to transcript ablation. Similarly, in the F2 56.9% of the CNV occurrences were predicted to lead to transcript amplification, followed by 21.6% to truncation of a feature, 9.8% to stop codon loss, 7.8% to ablation of the transcript and finally 3.9% to feature elongation. Because CNV events can be originated from the activity of transposable elements (TE), and metabolic challenges has been linked to dysregulation of TE, we investigated if there was an unusual enrichment of repetitive elements (RE) in the CNV regions by performing a permutation test on the levels of abundance of the annotated RE in the CNV regions. Indeed, significant enrichment of LINEs and LTR in four of the identified CNV events was observed (Supplementary Table 2), which were statistically significant when the levels of enrichment were compared between ON and CT (Supplementary Figure 4 and 5).

### Impact of metabolic challenge across 3 generations on the sperm epigenome

To determine if the multigenerational metabolic challenge impacts the sperm methylome of the mice, we performed GBS-MeDIP. To determine if the treatment affected overall methylation patterns, PCA on the methylated windows identified from the GBS-MeDIP libraries was performed. No clear separation in the PCA was observed (Figure 5), which indicates that the treatment did not pose a big impact on the overall methylome. This observation was further confirmed by the fact that only family belonging is correlated with PC2, and neither variable was correlated with the PC1 in the analysis of correlation between PCs and treatment/family belonging (Table 2).

**Figure 5.**
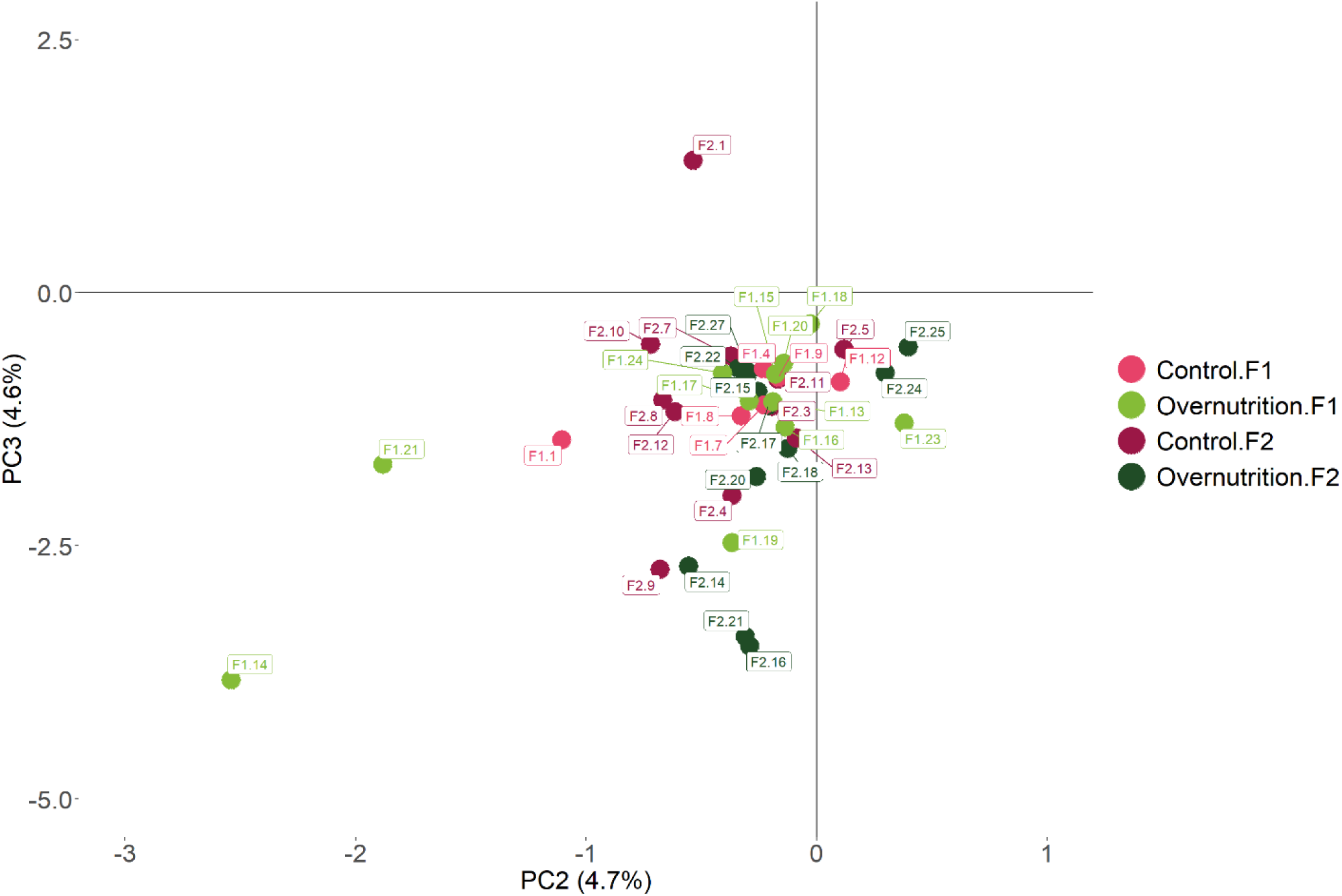
Principal component analysis (PCA) on the methylation counts from the GBS-MeDIP data. The individuals do not cluster by treatment, which is concordant with not being correlated with any PC.

**Table 2.**
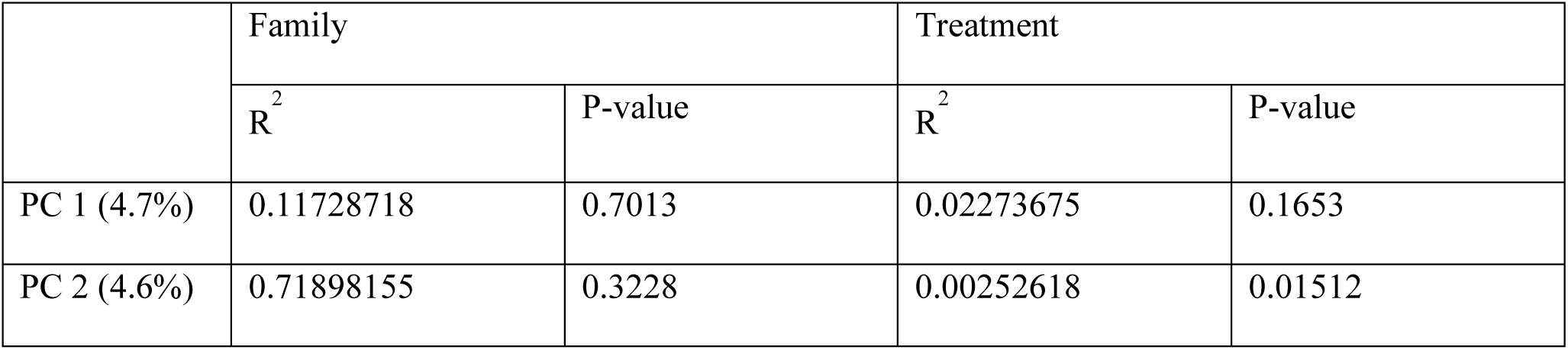
Association between treatment and families to the principal components of the PCA made with the epigenomic data. The data did not follow normality, Kruskal-Wallis test was used to assess the statistical significance of the correlation between the principal components and the family/treatment group.

|  | Family |  | Treatment |  |
| --- | --- | --- | --- | --- |
|  | R<sup>2</sup> | P-value | R<sup>2</sup> | P-value |
| PC 1 (4.7%) | 0.11728718 | 0.7013 | 0.02273675 | 0.1653 |
| PC 2 (4.6%) | 0.71898155 | 0.3228 | 0.00252618 | 0.01512 |

To explore more subtle changes that might not appear in the PCA, we analyzed the different density profiles of the methylation signals of ON and CT across the whole genome (Supplementary Figure 3). While the overall density profile was highly similar, subtle variations could be observed, suggesting potentially underlying changes that might converge at a higher level, such as biological pathways. To investigate this, we performed pathway enrichment analysis on genes annotated to windows from the GBS-MeDIP libraries containing reads that had a minimum length of 100 bps in each group and each generation. We identified pathways with lower FDR in the CT group and higher FDR or complete absence of the pathway in the ON group (Figure 6). This indicates weaker statistical evidence for enrichment of methylation-associated genes in these pathways in the ON group compared to the CT group.

**Figure 6.**
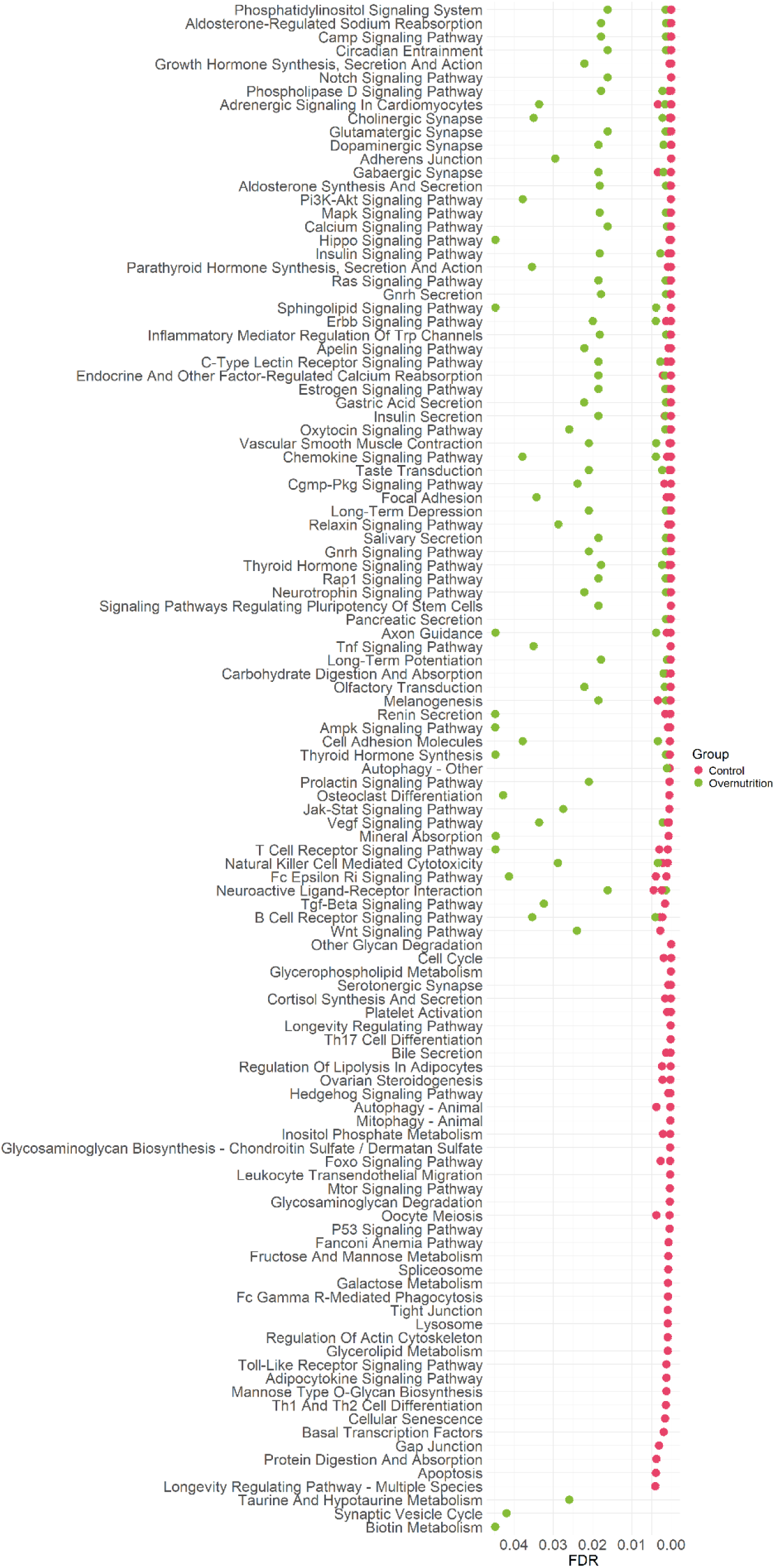
Significant pathways from the pathway enrichment analysis found in KEGG. The pathway enrichment was performed on genes annotated to windows containing reads that passed a minimum length of 100 bps in each group and each generation. Only the significant pathways are depicted. An the absense of signals in the ON group in same pathways as in CT is observed. Especially relevant are the absence of signal in metabolic pathways related to polysaccharides, lipolysis in adipocytes, and key developmental pathways.

Due to the enrichment of LINEs and LTR in the identified CNVs, we sought to explore potential relationships between DNA methylation and RE. To address this, we first examined the general distribution of REs in the GBS-MeDIP libraries, comparing the proportions of annotated transposable elements (TEs) between the CT and ON groups. No significant differences were found for any type of RE in the overall level of abundance (Figure 7). Expectedly, an enrichment of REs was observed in GBS-MeDIP libraries when comparing to GBS libraries. Subsequently, we compared DNA methylation levels at REs between groups. Methylation is maintained across both groups (ON and CT) and generations (Supplementary Figure 6) in a core of RE. However, in LINE and LTR elements, ON groups showed differential methylation levels in both generations (Supplementary Tables 3 and 4). This data suggests that the CNV events observed in the ON individuals might be due to dysregulation in the methylation patterns of RE in the germline genome.

**Figure 7.**
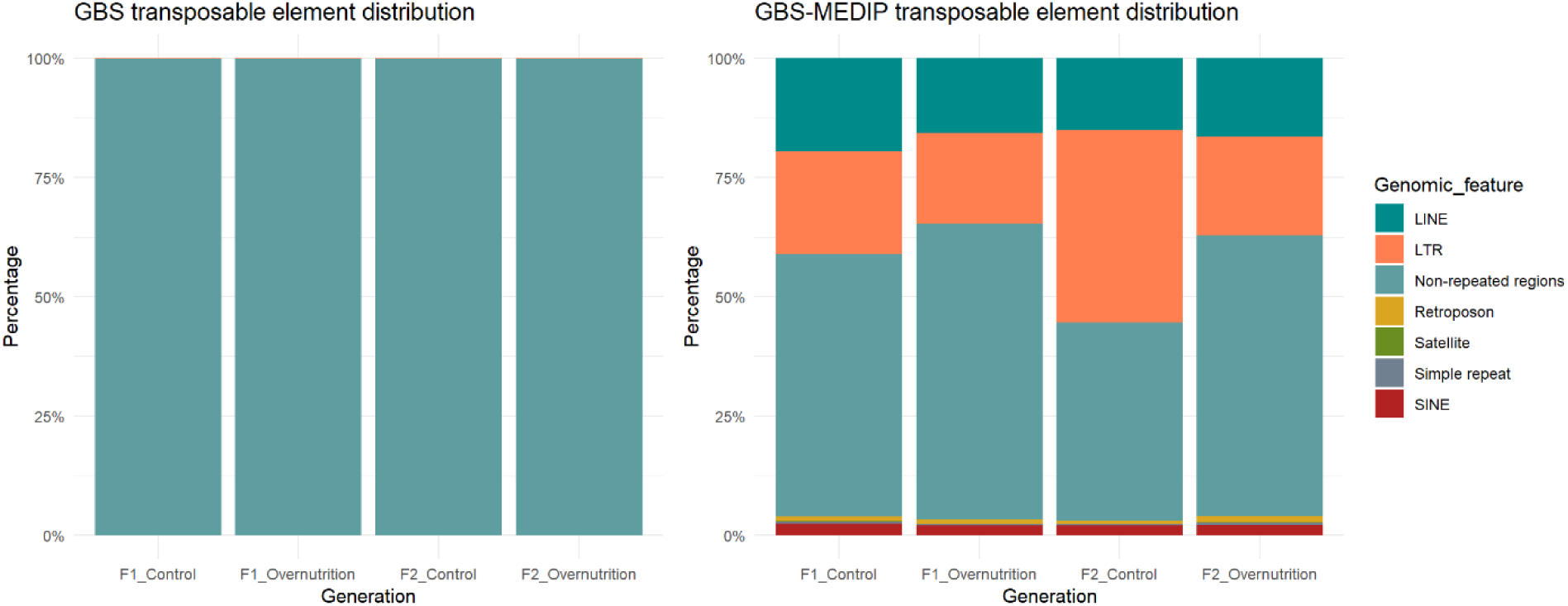
Bar plot featuring the percentage of repeated elements covered by the GBS and GBS-MeDIP library, respectively. Due to the process of immunoprecipitation, combined with the PtsI cleave site, which does not have bias against any genome feature, there is an enrichment in repeated elements due to their high levels of methylation. Although there are no significant differences between overall representation of any type of repeated element in any group.

Having observed subtle general differences in the methylome of the ON, we then performed differential methylation analysis to uncover potential regional epigenetic differences between ON and CT, two different types of analysis were done. To understand how the challenge affected the individuals, intragenerational comparison (CT vs ON) was performed in each generation. The second analysis was an intergenerational comparison, in between groups (F1 vs F2). This stems from the fact that we have several generations where the challenge is maintained, thus we investigated if new differences could arise across generations.

When performing the intragenerational comparison, we identified 5 DMRs in the F2 generation (Figure 8), 2 of them in the genes coding for *Fhod3 and Etl4,* one DMR was identified inside the satellite GSAT_MM, and the last two were found in LINEs and in SINEs. In the latter, the DMRs were hypomethylated in ON, while DMRs associated with *Fhod3 and Etl4* were hypermethylated in the same group. No DMRs were found in F1. However, having the advantage of data from multiple generations, we investigated the behaviour of the DMRs identified in the intragenerational analysis in F2 also in F1, although there was not an apparent pattern.

**Figure 8.**
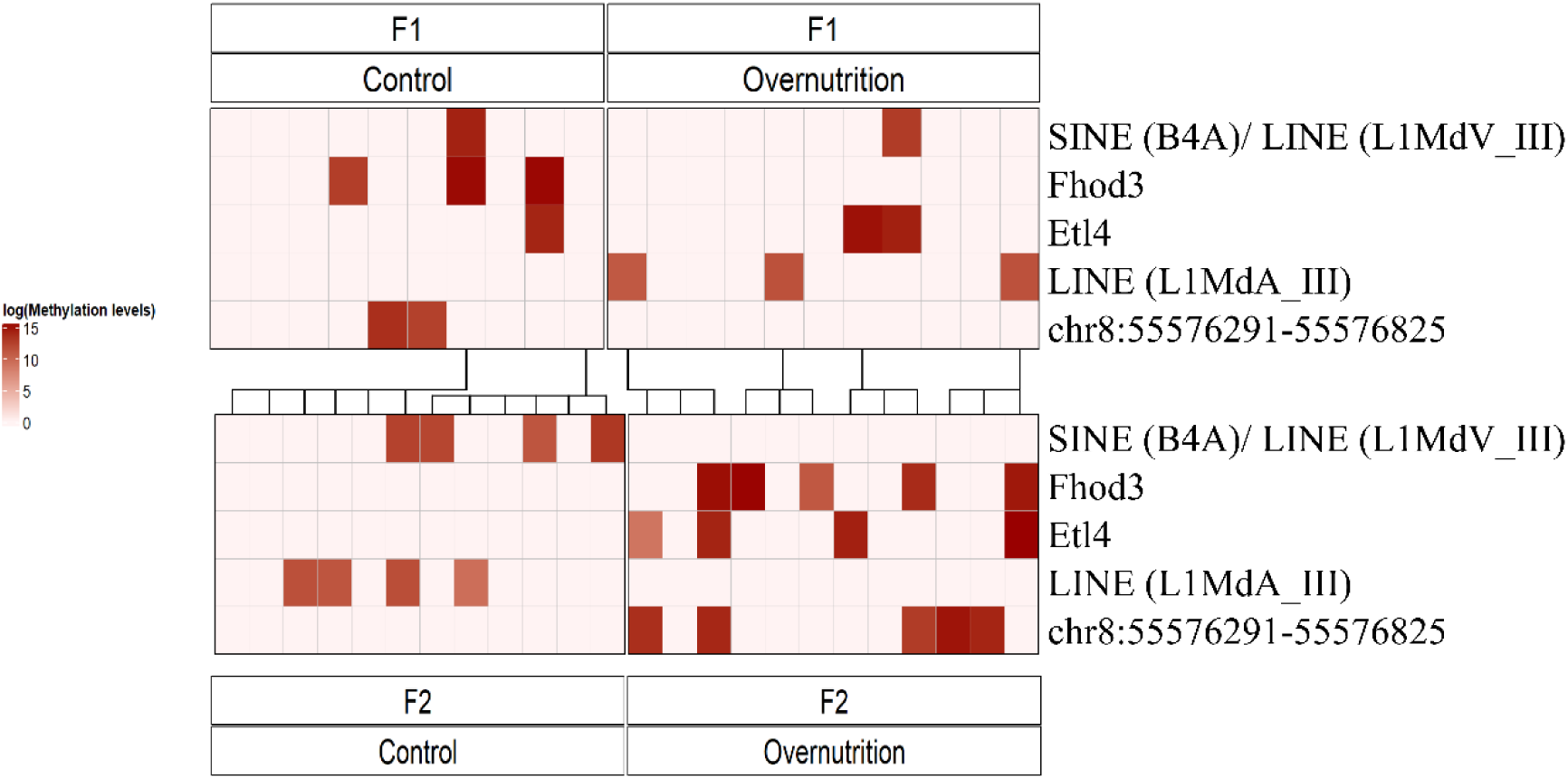
Intragenerational Differential Methylated Regions (DMR) found in the F2 generations. The level of methylation is in logarithmic scale to better visualize the changes. Each father giving offspring is connect to their descendants with black lines.

As the metabolic challenge was repeated in each generation, we also tested the possibility of having multiple hits in the different generations by performing an intergenerational comparison. In the ON, we found five DMRs, including two that overlapped with the intragenerational DMRs (Figure 9). These overlapping DMRs spanned CpG islands associated with the same satellite type, GSAT_MM, although they occurred on chromosomes 2 and 9. In contrast, no DMRs were detected in the CT group when comparing F1 and F2.

**Figure 9.**
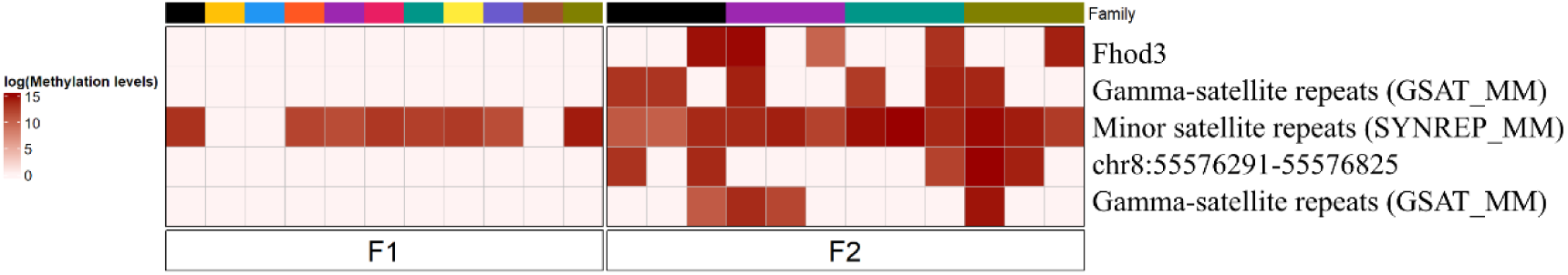
Intergenerational Differentially Methylated Regions (DMRs) for the ON comparing F1 vs F2 in order to observe different hits of the metabolic challenge in the different generations. The level of methylation is in logarithmic scale to better visualize changes. Each family is colour coded, we can observe fathers from F1 and offspring from F2.

### Interaction between DMRs and genetic changes

Because methylated cytosine is twelve times more likely to spontaneously transition to thymine, epimutations in the germline can be a source of new SNPs in further generations if the epimutation resist epigenetic reprogramming. To investigate the relation between methylation levels and SNPs, we tracked families across two generations (fathers of F1 and offspring of F2), focusing on the emergence of novel SNPs in offspring in relation to the methylation levels in fathers. In the CT we observed a slight positive correlation, having offspring with a higher SNP emergence in the same fragment the higher the methylation signal was in the parent. However, in the ON we observed a shift towards fewer SNPs emerging in the offspring in the same fragment the father had higher levels of methylation (Figure 10).

**Figure 10.**
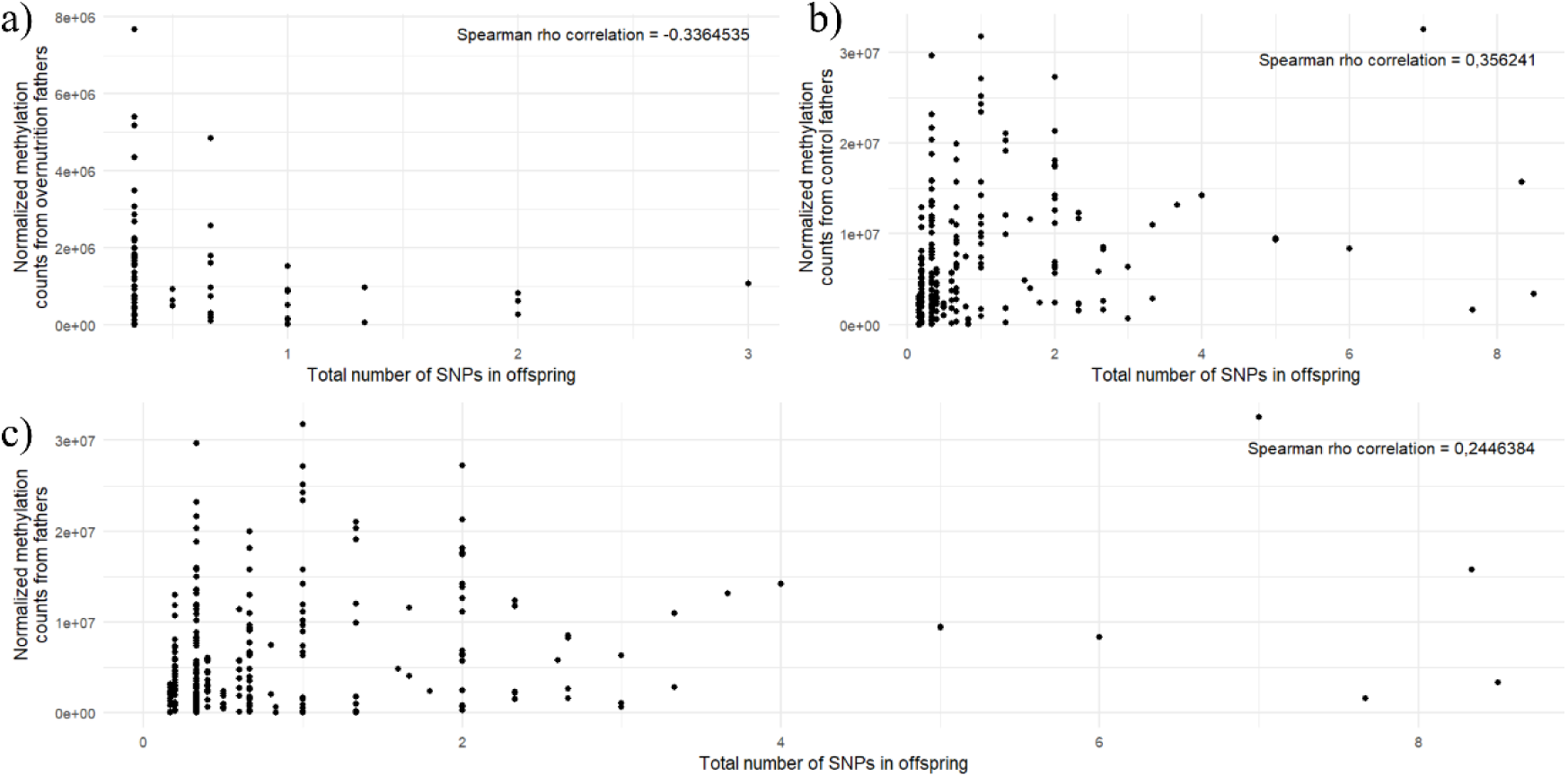
Representation of the correlation between novel SNPs in a) overnutrition group, b) control group and c) total. It is also attached the rho value from the spearman rank correlation. We see a weak negative trend on the ON and the opposite trend for the CT. When plotting together we see a weaker positive trend.

In addition, CNV can be produced by dysregulation of RE via loss of methylation. Therefore, we also investigated potential relations between methylation and CNVs, by examining the correlation between CNVs in offspring and methylation levels in fathers. We found that in the ON, none of the methylated windows in fathers corresponded to any CNV events, preventing any correlation analysis.

## Discussion

In this study we investigated in mice whether a multigenerational metabolic challenge affects the germline genome and methylome at different levels, and whether any interaction between the genome and the epigenome across generations exists. To assess whether the metabolic challenge affected the genome, we focused on SNP and CNV events to determine how the ON germline genome differed from the CT. Our PCA SNP-based analysis showed that the metabolic challenge clustered together unrelated families on the ON group, while on the CT group only family related individuals clustered together. This finding was further reinforced by the observation of a similar clustering pattern of unrelated families from the ON group on a hierarchical tree. Together these results show an effect of the treatment on the genetic variability of individuals. We also found that the treatment was associated with 4.6% of the genomic variance observed between CT and ON. Similar SNP clustering has been previously reported after environmental exposures. For example, exposure of populations of the Ribwort Plantain (*Plantago lanceolata*) to high CO_2_ levels over several generations contributed to ∼ 2% of the genetic variance seen between the exposed and the control populations (Saban et al., 2020). Animal studies have shown genomic diversification in relation to heavy metal exposure, where local populations exposed to lead and arsenic have genetically diverged on several loci over the course of multiple generations from unpolluted reference sites (Quina et al., 2021). In humans, studies on genotype-by-environment interactions show early-life environmental quality being linked to changes in population genetic makeup for reproductive timing (Mawass et al., 2022). These studies, however, were conducted with no information on number of generations, and without tracking families, which prevents the identification of genomic variation across generations, a key novelty of our study.

Beyond SNP-level genetic variation, we also investigated structural genomic variation. Because this study interrogated a reduced representation of the genome, we constructed a model with CT data that served as the template for calling CNV events on the ON individuals. We observed the emergence of CNV events in F1 ON individuals, and particularly in the case of deletions, we observed maintenance in F2. Interestingly, if the CNV event in the F1 was a duplication, we observed expansion of this event in the F2. These findings support a potential role for CNVs in genetic divergence driven by environmental exposures. Traditionally, CNV events have been investigated in humans or livestock in the context of their association to either illnesses or traits rather than related to their potential involvement in genomic diversification (Craddock et al., 2010); (Li et al., 2020); (Pös et al., 2021). However, there have been reports in humans and animals on the emergence of CNV in relation to exposures. In humans, for example, the emergence of novel CNV events has been reported in the germline offspring after exposure of both parents to caesium (Costa et al., 2018). In rats, CNVs events increased after a gestational exposure to vinclozolin, from the F1 to the F3 unexposed generations (Skinner et al., 2015).

The activity of TEs have been described as a source of structural genetic variation in animals that can influence the emergence of CNVs (Morgan et al., 1999); (Balachandran et al., 2022). Due to this potential, we examined the possibility that the identified CNVs in our study could relate to RE activity. We found a significant enrichment of the TEs LINEs and LTR inside the boundaries of the CNV events identified in the ON group, suggesting these CNVs could have emerged from TE activity. A previous study has shown correlations between LINEs and CNV emergence in the context of speciation related to chicken domestication (Pértille et al., 2019).

Having established that the metabolic challenge left measurable genomic impacts, we next investigated its epigenomic consequences, based on the bulk of literature showing germ line methylation effects of early life exposures (Stenz et al., 2018). Particularly, obesity and related metabolic diseases are known to promote epigenetic changes (Nätt & Öst, 2020) and they have been linked to changes in sperm DNA methylation in one generation (Jönsson et al., 2025) and across generations (Ben Maamar et al., 2023). Based on PCA clustering, we did not observe clusterization of ON or CT individuals, which is a different pattern compared to the SNP-based clustering performed on the genetic data. However, pathway enrichment analysis performed on the annotated genes related to methylation reads showed weaker statistical evidence for enrichment of these genes in the same pathways between the ON and the CT groups. Interestingly, some pathways are even absent in the ON group. This indicates methylation-dependent dysregulation of pathways in the ON group in both generations. Among the most affected, are pathways related to the metabolism of various polysaccharides (mannose, galactose, fructose), lipolysis in adipocytes, the P53 pathway, a crucial cell cycle check point protein related to apoptosis and cancer (Kuerbitz et al., 1992), and key developmental pathways such as mTOR and hedgehog. The mTOR pathway has been associated with insulin resistance (Zou et al., 2020), while the hedgehog pathway has been associated with diabetes and obesity (Garg et al., 2022). Therefore, despite a lack of global methylation effects triggered by the multigenerational early life metabolic disruption, we identify functional effects related to methylation, we could lead to specific methylation changes.

We then investigated whether different generations of ON harboured new germline DMRs. For this, we performed the frequently employed intragenerational methylomic comparisons between experimental treatments and additionally, we performed intergenerational comparisons within treatments between individuals of the F2 and F1 generations. In the intragenerational comparison, we did not identify sperm DMRs between ON and CT animals in the F1 generation, but we identified five DMRs in the F2 generation, two of them in gene bodies and three related to RE such as gamma-satellites repeats, LINEs and SINEs. One of the gene-body DMR was found in *Etl4*, which is related to regulation of the cell energy metabolism (Zhang et al., 2021), while the second was in *Fhod3*, related to the assembly of thin actin filaments and myofibrillogenesis in cardiomyocytes, helping to stabilize cardiac sarcomeres and having an active role in cardiac metabolism (Ochoa Juan et al., 2018). With the intergenerational comparison we found five DMRs, three of which were also observed in the intragenerational comparison. Interestingly, one DMR mapped again to *Fhod3,* and the others were related to the same gamma-satellites repeats, reinforcing the idea that the multigenerational metabolic challenge affected RE methylation. The finding of a DMR in *Fhod3* in both comparisons make this gene of special interest. *Fhod3* has been shown to be associated with thyroid cancer when hypomethylated and overexpressed (Chai et al., 2016).

Overall, with the methylomic analyses we did not identify PCA clustering of the individuals in the different experimental groups, as expected, since the impact of the treatment is limited to metabolic pathways and not the overall system. In later analysis, we identified specific DNA methylation changes related to metabolism, and mostly mapping to TEs, shedding light both on the functional consequences and mechanism related to these perturbations. Our findings are concordant with previous studies showing DNA methylation might not be the primary mechanism through which metabolic challenge is transmitted (Ribó et al., 2025); (Ribas-Aulinas et al., 2023) and are modest in provoking methylome changes, typically ranging from 10-20% of the changes observed (Jeremy M. et al., 2015); (Nätt & Öst, 2020). It is important to note that our analysis was on a reduced fraction of the genome, limiting our ability to identify all possible methylation changes across the genome.

Having examined the genetic and methylomic effects separately, we investigated possible connections between these levels. While in the CT group we found a positive correlation between methylation levels in the sperm of fathers and the emergence of SNPs in their offspring, we observed an inverse correlation in the ON group. Our findings in the CT group are concordant with the well-known increased mutation rate observed in methylated cytosines (Huttley, 2004); (Ying & Huttley, 2011). However, while the inverse correlation observed in the ON group was unexpected, one possible interpretation is that, under the metabolic challenge, methylated CpGs might be less prone to mutations. Although this could sound counterintuitive, this possibility is consistent with reports showing a protective role for CpG methylation against chemically induced mutations or damage. For example, the conceptual core of the bisulphite sequencing technique is that methylated CpGs are protected from mutation induced by sodium bisulphite (Clark et al., 2006). More recently, a reduced susceptibility of CpGs to oxidation-induced damage has been reported (Takhaveev et al., 2026). Although methylation was not assessed in the study, the reduced oxidation of CpGs could be partially related to DNA methylation. Taken together, these observations raise the possibility that two seemingly opposite processes may be operating simultaneously. Under baseline conditions (e.g., in the CT group), CpG methylation may bias the emergence of SNPs, consistent with its known higher mutational rates. However, under metabolic challenge (ON group), methylated CpGs may be less affected by damage-associated mutagenesis, a process potentially mediated by oxidative stress.

The second integration aimed to investigate potential relationship between structural genetic variations and DNA methylation Previous work already suggest a relationship between CNV events and DNA methylation in the sperm. transgenerational CNV events in sperm after a gestational exposures of rats to vinclozolin, which increased from the gestationally exposed F1 generation to the unexposed F3 generation (Skinner et al., 2015), as well as sperm methylomic changes in the same model observed in line elements in the F3 generation (Guerrero-Bosagna et al., 2010). Our current results showing enrichment of LINEs and LTR elements in the CNVs of the ON group, and the emergence of DMRs in both the intra and intergenerational comparisons mapping to RE, particularly to LINEs, point to the same direction. Further supporting this idea, we observed differential methylation in the LINEs for both F1 and F2 generations between ON and CT in the DMRs. This is supported by previous results of DMRs which were related to RE and CpG islands in satellites, with hypomethylation in F1 followed by a new methylation wave in F2. Even though these results are correlation analysis, we are indicating for future reference that REs might be the link between epimutations carried in the germline and CNV emergence in subsequent generations.

Our study demonstrates how a multigenerational metabolic challenge affects the sperm genome and methylome across generations. We observed that, compared to the CT group, the ON group exhibits constrained genetic variation at the level of SNPs, but higher genome instability related to CNV events. The latter seems to be arising from the dysregulation of RE methylation. On the methylomic part, although we observed limited regional changes, we identified several pathways potentially affected by combined small methylomic effects. More affected pathways are involved in polysaccharide metabolism, cell cycle check point regulation, and key developmental pathways. Additionally, it seems that methylation changes identified affect the activity of repeat and transposable elements across generations. Finally, we identify a gene altered across generations via two different comparisons, namely *Fhod3*, involved in thyroid cancer when hypomethylated.

## Acknowledgements

This study was funded by Templeton foundation under grant number 62167. Special thanks to NAISS, the National Academic Infrastructure for Super-computing in Sweden for the access to the UPPMAX server. The authors want to acknowledge the help from Judith Cebria in performing animal breeding in the animal facility.

## Data availability

The data underlying this article are available in ENA project accession number: PRJEB105468 for GBS and GBS-MeDIP data. Scripts used in this article are in https://github.com/Violeta-de-Anca/Evolution_of_SNP_emerging_in_relation_to_methylation_fragments_in_an_obesogenic_environment.git

