## Supplementary Information 1 for "Germline genomic and methylomic dynamics following three generations of early-life metabolic challenges"

### Supplementary material

#### Site Frequency spectrum

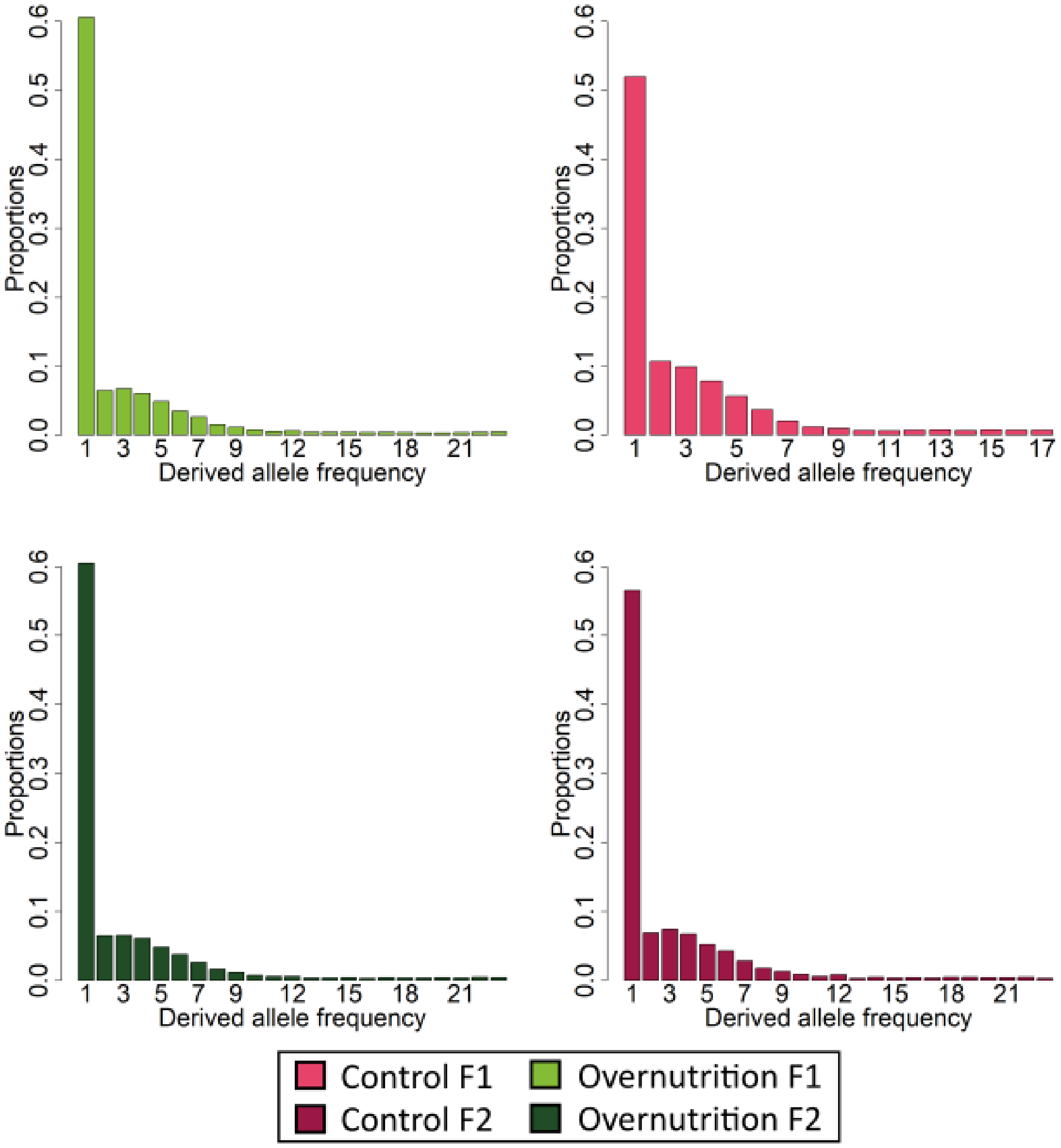


*Supplementary Figure 1. Site Frequency Spectrum for overnutrition group and control in F1 and F2 generations. In shades of pink, we find the two different generations of control. In shades of green we find the two generations belonging to the overnutrition group. The upper graphs are the F1 generations and the graphs below the F2.*

### PCA from SNP on individuals

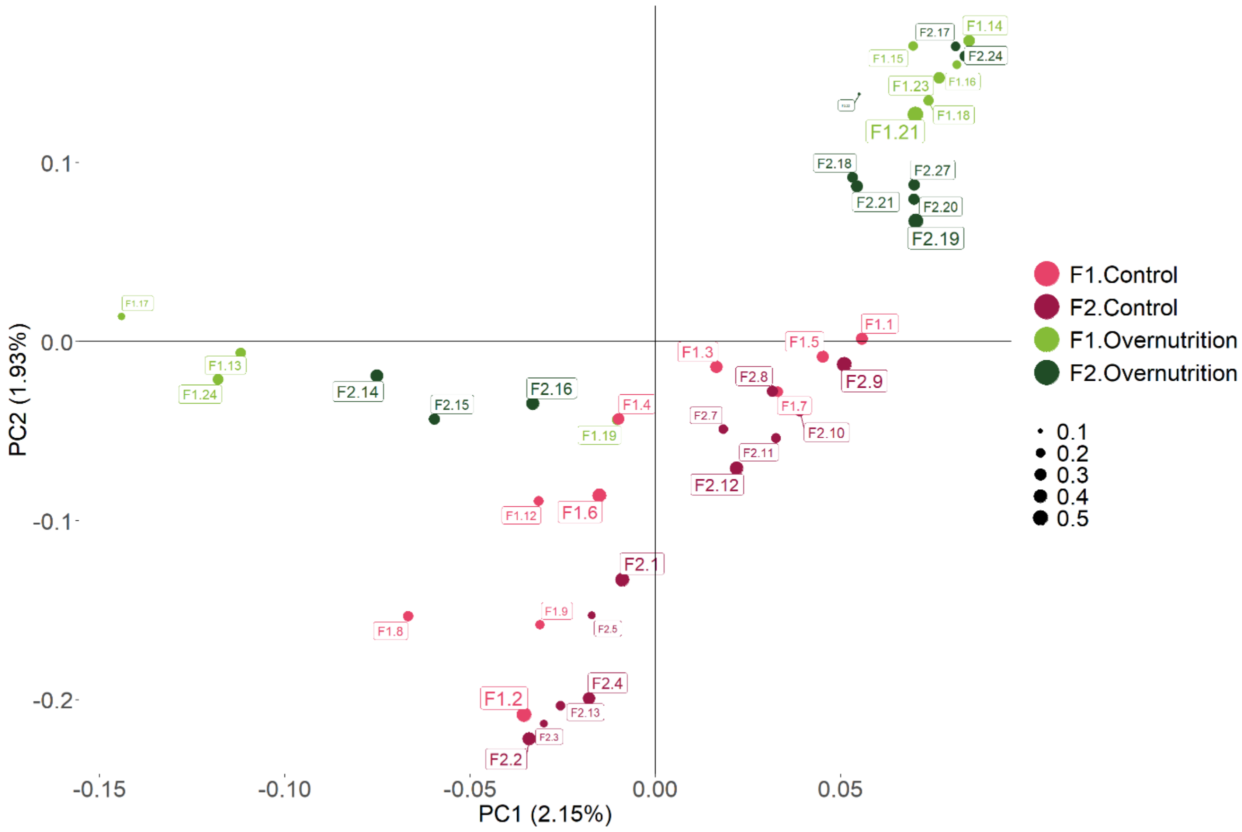


*Supplementary Figure 2. Principal Component Analysis using PLINK, where also sequencing depth of each individual is taken into account depicted as the size of the circles. The different clusters are composed of several individuals with different coverage, meaning the clusterization is due to different factors such as family or group belonging.*

### Quality control of the GBS and GBS-MeDIP

*Supplementary Table 1. Quality control was performed over each individual for their GBS and GBS-MeDIP library.*

| | **GBS** | | | | **GBS-MeDIP** | | | | | | | |
|---|---|---|---|---|---|---|---|---|---|---|---|---|
| Sample Name | M Reads uniquely Mapped | Number SNPs before | Number of SNPs after | Coverage | M Reads Uniquely Mapped | Number of peaks in MACS3 | M Reads MQ >10 Mapped | Ratio mapped GBS/mapped GBS-MeDIP | Coverage | Number of peaks in MACS3 | Windows with GBS info | Retention of peaks |

| | | | | | | | | | | | | |
|---|---|---|---|---|---|---|---|---|---|---|---|---|
| | | filtering | filtering | | | | | | | with MQ >10 | | |
| F1_1 | 9532346 | 71325 | 56979 | 0,48 | 799434 | 981 | 882868 | 11,92 | 0,04 | 753 | 239 | 31,74 |
| F1_10 | 7875170 | 53646 | 42945 | 0,41 | 882868 | 1609 | 419284 | 8,92 | 0,05 | 1122 | 433 | 38,59 |
| F1_11 | 2949167 | 43285 | 31584 | 0,15 | 419284 | 823 | 799434 | 7,03 | 0,02 | 531 | 146 | 27,50 |
| F1_2 | 11993655 | 64899 | 52347 | 0,62 | 1433180 | 2352 | 1433180 | 8,37 | 0,07 | 1591 | 689 | 43,31 |
| F1_3 | 6813498 | 52597 | 41669 | 0,35 | 668024 | 1248 | 668024 | 10,20 | 0,03 | 857 | 382 | 44,57 |
| F1_4 | 6579685 | 57683 | 44977 | 0,34 | 735610 | 1188 | 735610 | 8,94 | 0,04 | 926 | 274 | 29,59 |
| F1_5 | 2789060 | 52468 | 36499 | 0,15 | 475478 | 864 | 475478 | 5,87 | 0,02 | 613 | 234 | 38,17 |
| F1_6 | 5521051 | 59498 | 45471 | 0,29 | 728864 | 1104 | 728864 | 7,57 | 0,04 | 866 | 348 | 40,18 |
| F1_7 | 8125653 | 56011 | 44477 | 0,42 | 859692 | 1465 | 859692 | 9,45 | 0,05 | 938 | 416 | 44,35 |
| F1_8 | 15796108 | 70511 | 57607 | 0,83 | 1126696 | 1712 | 1126696 | 14,02 | 0,06 | 1249 | 520 | 41,63 |
| F1_9 | 3819893 | 57926 | 44293 | 0,20 | 664342 | 1309 | 664342 | 5,75 | 0,03 | 965 | 185 | 19,17 |
| F2_1 | 5651945 | 54757 | 42968 | 0,30 | 772034 | 1248 | 434946 | 7,32 | 0,04 | 977 | 265 | 27,12 |
| F2_12 | 4099147 | 41508 | 31274 | 0,21 | 434946 | 704 | 339730 | 9,42 | 0,02 | 556 | 129 | 23,20 |

| | | | | | | | | | | | | |
|---|---|---|---|---|---|---|---|---|---|---|---|---|
| F2_13 | 4591681 | 43252 | 32854 | 0,24 | 339730 | 658 | 538870 | 13,52 | 0,02 | 464 | 116 | 25,00 |
| F2_14 | 5413092 | 47075 | 36098 | 0,28 | 538870 | 1076 | 301630 | 10,05 | 0,03 | 797 | 262 | 32,87 |
| F2_15 | 3916883 | 43107 | 31697 | 0,20 | 301630 | 716 | 432692 | 12,99 | 0,02 | 513 | 84 | 16,37 |
| F2_16 | 3401582 | 45681 | 32215 | 0,18 | 432692 | 691 | 400382 | 7,86 | 0,02 | 560 | 165 | 29,46 |
| F2_17 | 2784087 | 42217 | 29451 | 0,14 | 400382 | 607 | 666280 | 6,95 | 0,02 | 491 | 149 | 30,35 |
| F2_18 | 4553612 | 53542 | 41150 | 0,24 | 666280 | 1138 | 453476 | 6,83 | 0,03 | 844 | 233 | 27,61 |
| F2_19 | 4925775 | 45537 | 35277 | 0,26 | 453476 | 731 | 772034 | 10,86 | 0,02 | 558 | 174 | 31,18 |
| F2_2 | 9266197 | 64205 | 52008 | 0,48 | 974618 | 1607 | 384766 | 9,51 | 0,05 | 1259 | 436 | 34,63 |
| F2_20 | 772118 | 29386 | 17416 | 0,04 | 384766 | 820 | 856032 | 2,01 | 0,02 | 617 | 40 | 6,48 |
| F2_21 | 9943463 | 59371 | 48388 | 0,52 | 856032 | 1487 | 498998 | 11,62 | 0,04 | 1029 | 341 | 33,14 |
| F2_23 | 5637068 | 51028 | 39803 | 0,29 | 498998 | 957 | 503880 | 11,30 | 0,03 | 674 | 193 | 28,64 |
| F2_24 | 4758245 | 48659 | 36788 | 0,25 | 503880 | 841 | 974618 | 9,44 | 0,03 | 595 | 144 | 24,20 |
| F2_3 | 6312623 | 55671 | 44120 | 0,33 | 737208 | 1462 | 737208 | 8,56 | 0,04 | 1010 | 375 | 37,13 |
| F2_4 | 5394814 | 47792 | 37116 | 0,28 | 470446 | 1011 | 470446 | 11,47 | 0,02 | 732 | 161 | 21,99 |
| F2_5 | 5994771 | 47490 | 36639 | 0,31 | 742518 | 1051 | 742518 | 8,07 | 0,04 | 876 | 329 | 37,56 |

| | | | | | | | | | | | | |
|---|---|---|---|---|---|---|---|---|---|---|---|---|
| F2_6 | 8066207 | 60566 | 48524 | 0,42 | 949780 | 1711 | 949780 | 8,49 | 0,05 | 1213 | 450 | 37,10 |
| F2_7 | 4538241 | 43478 | 33288 | 0,24 | 361830 | 756 | 361830 | 12,54 | 0,02 | 566 | 111 | 19,61 |
| F2_8 | 4479488 | 46816 | 35733 | 0,24 | 392776 | 874 | 392776 | 11,40 | 0,02 | 604 | 137 | 22,68 |
| F2_9 | 3578506 | 40802 | 30305 | 0,19 | 367038 | 624 | 367038 | 9,75 | 0,02 | 435 | 110 | 25,29 |
| F3_1 | 8099423 | 58232 | 46190 | 0,42 | 785698 | 1143 | 488008 | 10,31 | 0,04 | 902 | 365 | 40,47 |
| F3_10 | 5537688 | 50900 | 39402 | 0,29 | 488008 | 968 | 292278 | 11,35 | 0,03 | 646 | 185 | 28,64 |
| F3_11 | 4172501 | 42802 | 32298 | 0,22 | 292278 | 642 | 599002 | 14,28 | 0,02 | 415 | 109 | 26,27 |
| F3_12 | 7819884 | 54072 | 43359 | 0,41 | 599002 | 1217 | 458022 | 13,05 | 0,03 | 855 | 209 | 24,44 |
| F3_13 | 3781192 | 44787 | 33143 | 0,20 | 458022 | 1004 | 574960 | 8,26 | 0,02 | 706 | 197 | 27,90 |
| F3_14 | 6842083 | 49798 | 39669 | 0,36 | 574960 | 1172 | 407534 | 11,90 | 0,03 | 799 | 294 | 36,80 |
| F3_15 | 5023711 | 45592 | 34935 | 0,26 | 407534 | 707 | 742954 | 12,33 | 0,02 | 507 | 144 | 28,40 |
| F3_16 | 7332678 | 53172 | 41956 | 0,38 | 742954 | 1114 | 413794 | 9,87 | 0,04 | 841 | 350 | 41,62 |
| F3_17 | 4023366 | 45118 | 33292 | 0,21 | 413794 | 880 | 450982 | 9,72 | 0,02 | 585 | 175 | 29,91 |
| F3_18 | 4768593 | 58517 | 43793 | 0,25 | 450982 | 944 | 881006 | 10,57 | 0,02 | 669 | 241 | 36,02 |
| F3_19 | 8934543 | 68926 | 55576 | 0,47 | 881006 | 1549 | 785698 | 10,14 | 0,05 | 1015 | 399 | 39,31 |

| | | | | | | | | | | | | |
|---|---|---|---|---|---|---|---|---|---|---|---|---|
| F3_2 | 7182664 | 59444 | 47315 | 0,37 | 831778 | 1375 | 609686 | 8,64 | 0,04 | 992 | 360 | 36,29 |
| F3_20 | 5080791 | 49903 | 38201 | 0,26 | 609686 | 1150 | 620400 | 8,33 | 0,03 | 788 | 256 | 32,49 |
| F3_21 | 5847632 | 52819 | 40959 | 0,30 | 620400 | 1205 | 275368 | 9,43 | 0,03 | 886 | 276 | 31,15 |
| F3_22 | 1783558 | 40536 | 25636 | 0,09 | 275368 | 487 | 510188 | 6,48 | 0,01 | 356 | 90 | 25,28 |
| F3_24 | 4665574 | 53970 | 40914 | 0,24 | 510188 | 846 | 438096 | 9,14 | 0,03 | 626 | 170 | 27,16 |
| F3_25 | 871396 | 45935 | 29890 | 0,05 | 438096 | 911 | 416536 | 1,99 | 0,02 | 706 | 82 | 11,61 |
| F3_27 | 5559676 | 50300 | 38301 | 0,29 | 416536 | 899 | 831778 | 13,35 | 0,02 | 597 | 148 | 24,79 |
| F3_3 | 2946439 | 41579 | 30521 | 0,15 | 327948 | 744 | 327948 | 8,98 | 0,02 | 519 | 142 | 27,36 |
| F3_4 | 6143055 | 48705 | 38248 | 0,32 | 583706 | 1027 | 583706 | 10,52 | 0,03 | 792 | 228 | 28,79 |
| F3_5 | 2876527 | 50844 | 38215 | 0,15 | 468928 | 1103 | 468928 | 6,13 | 0,02 | 790 | 143 | 18,10 |
| F3_7 | 3905425 | 42004 | 31114 | 0,20 | 359092 | 650 | 359092 | 10,88 | 0,02 | 515 | 98 | 19,03 |
| F3_8 | 5330231 | 50262 | 38746 | 0,28 | 406340 | 719 | 406340 | 13,12 | 0,02 | 493 | 196 | 39,76 |
| F3_9 | 8823174 | 61868 | 49782 | 0,46 | 714974 | 1200 | 714974 | 12,34 | 0,04 | 861 | 299 | 34,73 |

14

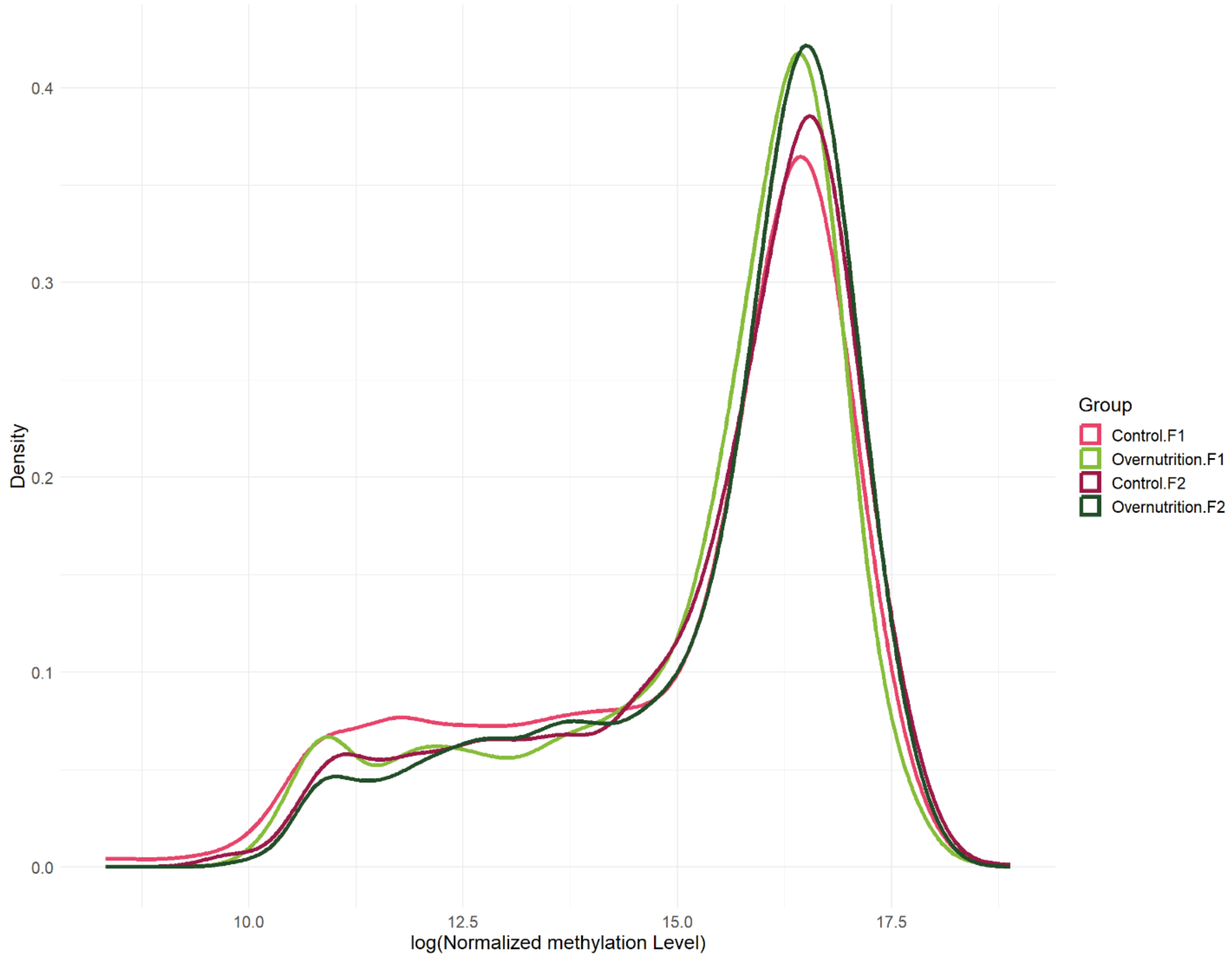


*Supplementary Figure 3. Density plots of the methylation levels. Control groups are shown in shades of pink and overnutrition groups in shades of green.*

### Repeated elements annotated in CNV events in ON

*Supplementary Table 2. Permutation test results of the enrichment of different types of RE in the CNV events detected in the ON group.*

| CNV coordinate | Type of RE significant | Experimental p-value |
|---|---|---|
| chr1:139085001-140139000 | LTR | 0.024 |
| chr17:33724001-34838000 | LTR | 0.030 |
| chr4:137356001-138024000 | LTR | 0.014 |

| | | |
|---|---|---|
| chr4:137356001-138024000 | LINE | 0.042 |
| chr8:70803001-71885000 | LTR | 0.024 |
| chr8:70803001-71885000 | LINE | 0.043 |

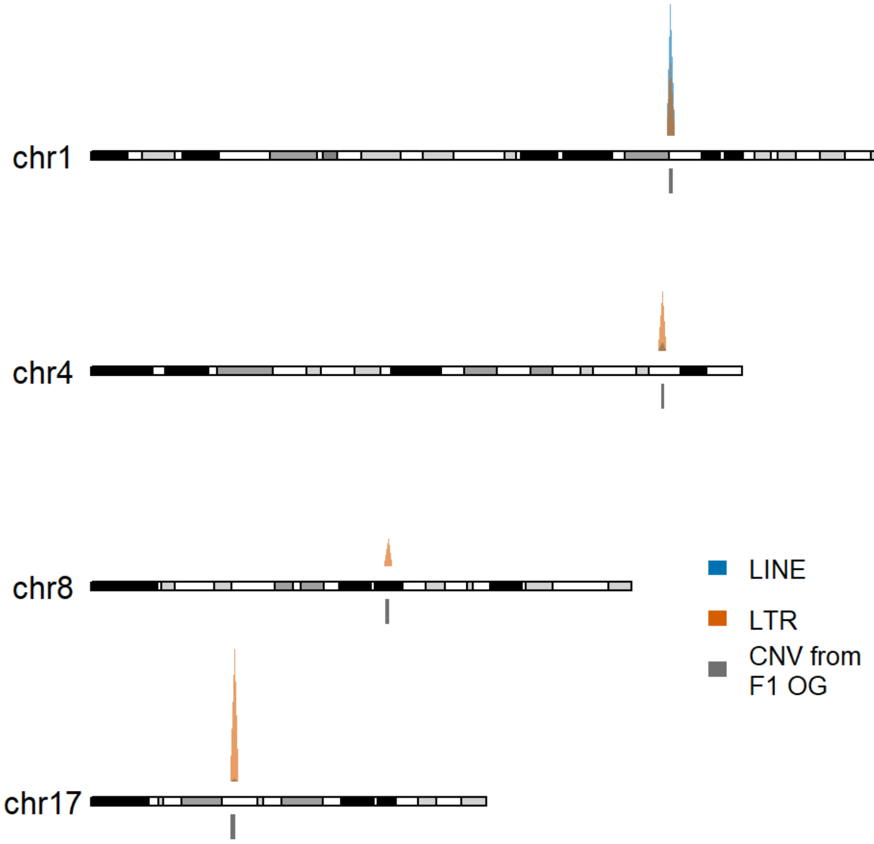


*Supplementary Figure 4. Annotation of repeated elements in the CNV events significantly differentially abundant identified in F1 generation in ON.*

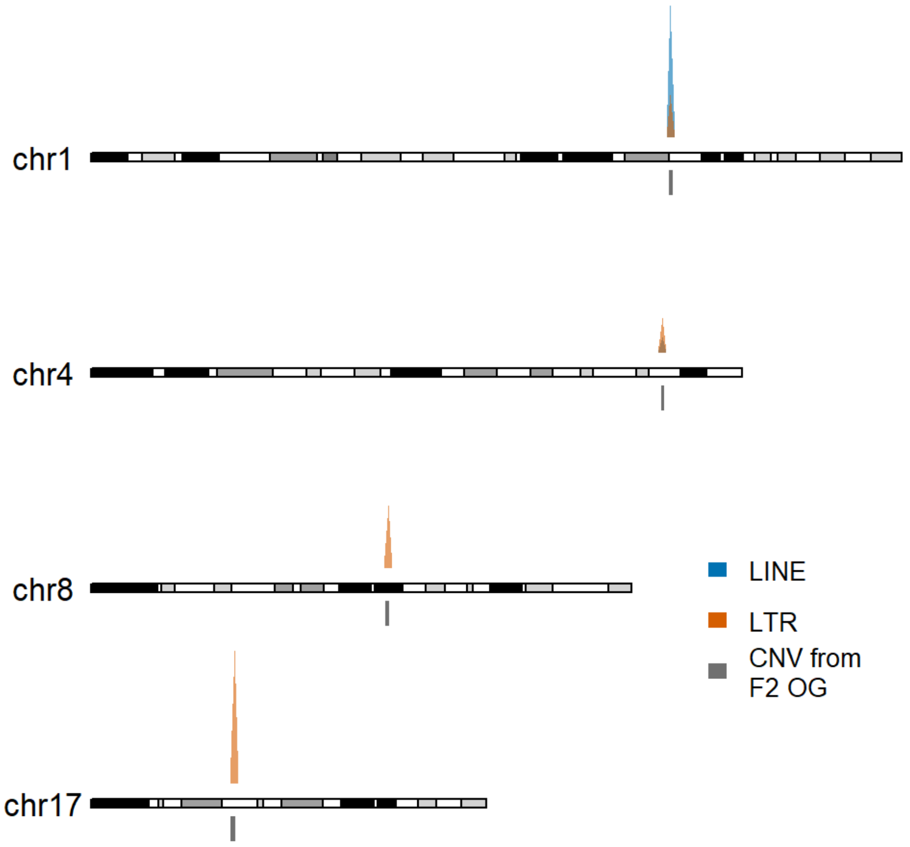


*Supplementary Figure 5. Annotation of repeated elements in the CNV events significantly differentially abundant identified in F2 generation in ON.*

### Repeated elements annotated in methylated windows

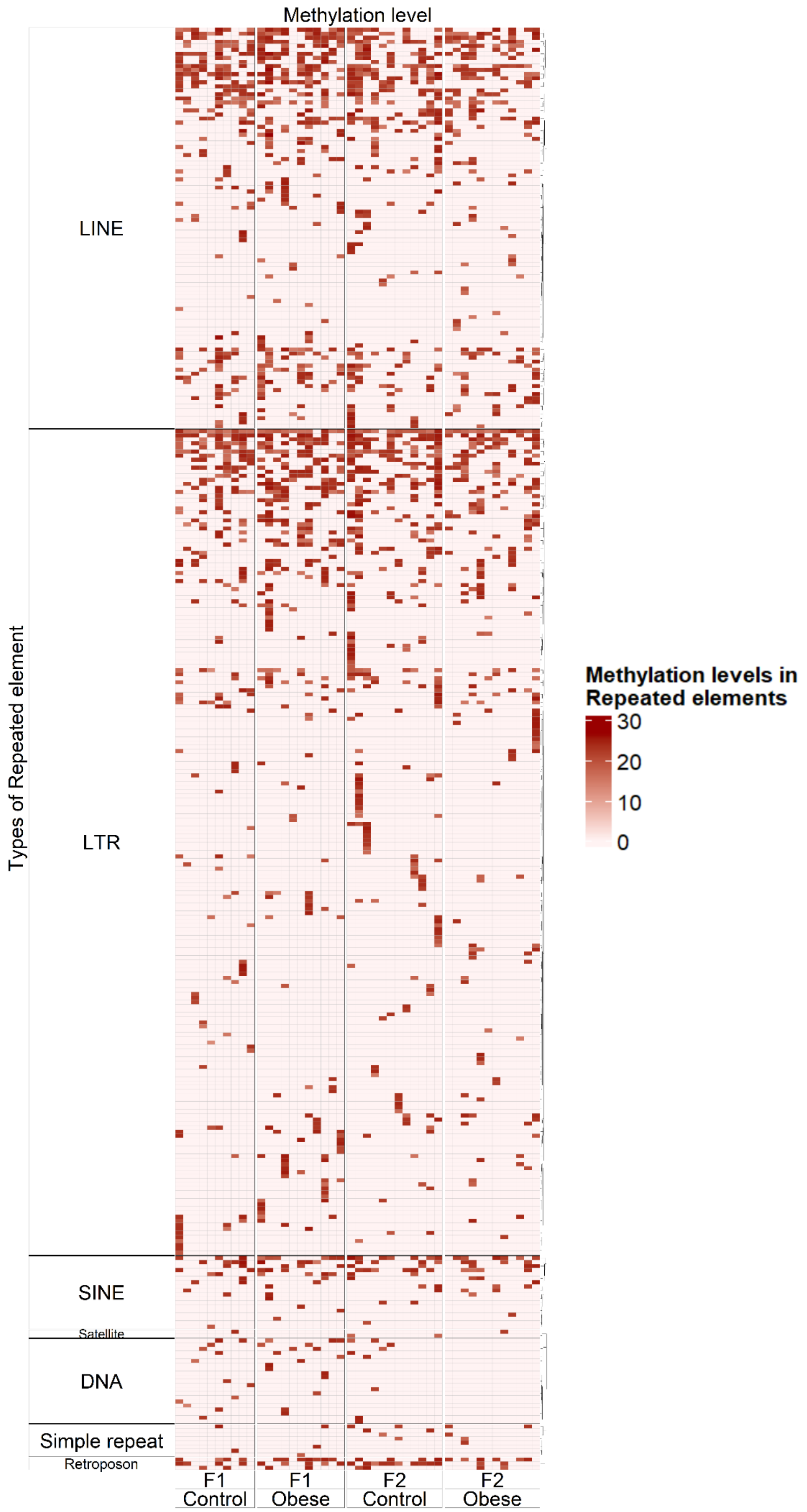

*Supplementary Figure 6. Annotation of repeated elements in methylated windows. Observe core repression of core repeated elements in all groups while other RE were dysregulated.*

### Statistical analysis of repeated elements annotated from GBS-MeDIP

*Supplementary Table 3. P-values of the identified RE in the methylome data comparing ON and CT of F1 generation. NA values are due to low individuals.*

| P-value | RE | General type of RE |
|---|---|---|
| 0.5 | B3 | SINE |
| 0.4 | B3A | SINE |
| 0.628571 | B4 | SINE |
| 1 | B4A | SINE |
| 0.28162 | IAPEz-int | LTR |
| 1 | IAPLTR1_Mm | LTR |
| 0.071429 | IAPLTR1a_Mm | LTR |
| 0.638889 | IAPLTR2_Mm | LTR |
| 0.8 | IAPLTR2a | LTR |
| 0.333333 | IAPLTR2a2_Mm | LTR |
| 1 | IAPLTR2b | LTR |
| **0.029526** | **L1Lx_I** | **LINE** |
| 0.8 | L1Lx_III | LINE |
| 1 | L1Lx_IV | LINE |
| 0.666667 | L1M2 | LINE |
| 0.857143 | L1M4 | LINE |
| 0.666667 | L1M5 | LINE |
| 1 | L1MA4A | LINE |

| | | |
|---|---|---|
| 1 | L1MA6 | LINE |
| 0.857143 | L1MB4 | LINE |
| 1 | L1MC1 | LINE |
| 1 | L1MD2 | LINE |
| NA | L1MDa | LINE |
| 0.666667 | L1MEc | LINE |
| 0.930736 | L1MdA_I | LINE |
| 0.523699 | L1MdA_II | LINE |
| 0.114286 | L1MdA_III | LINE |
| 0.2 | L1MdA_V | LINE |
| 0.730159 | L1MdA_VI | LINE |
| 0.904762 | L1MdA_VII | LINE |
| 0.8 | L1MdF_I | LINE |
| 0.901515 | L1MdF_III | LINE |
| 0.516667 | L1MdF_IV | LINE |
| 0.087912 | L1MdF_V | LINE |
| 0.533333 | L1MdFanc_I | LINE |
| 0.857143 | L1MdFanc_II | LINE |
| 0.571429 | L1MdGf_II | LINE |
| 0.261905 | L1MdMus_I | LINE |
| NA | L1MdMus_II | LINE |
| 1 | L1MdV_II | LINE |
| 0.8 | L1_Mur1 | LINE |
| 0.547619 | L1_Mur2 | LINE |
| 0.8 | L1_Rod | LINE |
| 1 | L2a | LINE |
| 0.666667 | L2b | LINE |

| | | |
|---|---|---|
| 1 | Lx | LINE |
| NA | Lx2B2 | LINE |
| NA | Lx3A | LINE |
| 1 | Lx3B | LINE |
| 0.666667 | Lx4A | LINE |
| 0.547619 | Lx5 | LINE |
| 1 | Lx5c | LINE |
| **0.042624** | **Lx6** | **LINE** |
| 0.555556 | Lx7 | LINE |
| 0.267677 | Lx8 | LINE |
| 1 | Lx8b | LINE |
| 0.392857 | Lx9 | LINE |
| 1 | MERVL-int | LTR |
| 0.8 | MERVL_2A-int | LTR |
| 0.7 | MMERVK10C-int | LTR |
| 0.8 | MMERVK9C_I | LTR |
| 0.8 | MMVL30-int | LTR |
| 0.428571 | MTA_Mm | LTR |
| NA | MTB | LTR |
| 1 | MTC | LTR |
| 0.309524 | MTD | LTR |
| 0.666667 | MTE-int | LTR |
| 0.4 | MTEa | LTR |
| 0.5 | MuRRS-int | LTR |
| 1 | ORR1A0 | LTR |
| 0.8 | ORR1A2 | LTR |
| 0.171429 | ORR1B1-int | LTR |

| | | |
|---|---|---|
| 0.666667 | ORR1B2 | LTR |
| NA | ORR1C1 | LTR |
| 0.666667 | ORR1C2 | LTR |
| 0.431818 | ORR1D1 | LTR |
| 0.666667 | ORR1D2 | LTR |
| 0.714286 | ORR1F | LTR |
| 0.5 | ORR1G | LTR |
| 1 | RLTR10 | LTR |
| 0.904762 | RLTR10-int | LTR |
| 0.114286 | RLTR20C1_MM | LTR |
| NA | RLTR40 | LTR |
| 0.547619 | RLTR4_MM-int | LTR |
| 0.057143 | RMER15 | LTR |
| 1 | RMER15-int | LTR |
| NA | RMER19B2 | LTR |
| 0.785714 | RMER1B | Retroposon |
| NA | RMER30 | DNA |
| 1 | RMER4B | LTR |
| 0.426167 | not_RE | . |
| 0.666667 | (GTGTGT)n | Simple_repeat |
| 0.666667 | B1F | SINE |
| 1 | B1_Mm | SINE |
| NA | ERV3-16A3_I | LTR |
| 1 | ERVB4_2-I_MM | LTR |
| 1 | ERVL-int | LTR |
| 1 | ETnERV-int | LTR |
| 0.666667 | ETnERV2-int | LTR |

| | | |
|---|---|---|
| NA | HERV16-int | LTR |
| 1 | IAP-d-int | LTR |
| 1 | IAPEY2_LTR | LTR |
| 0.082984 | IAPEY3-int | LTR |
| 1 | IAPEY5_I | LTR |
| NA | IAPEy-int | LTR |
| 0.25 | IAPLTR3-int | LTR |
| 0.666667 | IAPLTR4_I | LTR |
| 0.666667 | ID_B1 | SINE |
| NA | L1M3 | LINE |
| NA | L1M3c | LINE |
| 1 | L1M4b | LINE |
| NA | L1M4c | LINE |
| 0.666667 | L1MA4 | LINE |
| NA | L1MA5 | LINE |
| 1 | L1MA9 | LINE |
| NA | L1MB3 | LINE |
| NA | L1MB5 | LINE |
| NA | L1MB7 | LINE |
| 0.666667 | L1MB8 | LINE |
| NA | L1MCa | LINE |
| 0.666667 | L1ME1 | LINE |
| NA | L1MEg | LINE |
| 1 | L1MdV_I | LINE |
| NA | L1MdV_III | LINE |
| 0.914286 | L1_Mur3 | LINE |
| 1 | L2 | LINE |

| | | |
|---|---|---|
| NA | L2c | LINE |
| 1 | L3 | LINE |
| 0.666667 | Lx2 | LINE |
| 0.666667 | Lx2A | LINE |
| NA | Lx2B | LINE |
| NA | Lx3_Mus | LINE |
| 1 | Lx4B | LINE |
| 1 | Lx5b | LINE |
| 1 | MER2 | DNA |
| 1 | MIRb | SINE |
| 1 | MIRc | SINE |
| NA | MLT-int | LTR |
| NA | MLT1A | LTR |
| NA | MLT1A0 | LTR |
| 1 | MLT1B | LTR |
| NA | MLT1D | LTR |
| 1 | MLT1H | LTR |
| NA | MLT1K | LTR |
| NA | MMERVK10D3_I | LTR |
| 0.857143 | MMERVK9E_I | LTR |
| 0.8 | MT2A | LTR |
| 0.666667 | MT2B1 | LTR |
| 1 | MT2B2 | LTR |
| NA | MTB_Mm | LTR |
| NA | MTC-int | LTR |
| 1 | MTE2a | LTR |
| 0.666667 | MTE2b | LTR |

| | | |
|---|---|---|
| NA | MTEb | LTR |
| 0.666667 | MYSERV-int | LTR |
| NA | MYSERV16_I | LTR |
| 0.666667 | MuRRS4-int | LTR |
| NA | MurERV4-int | LTR |
| NA | ORR1A1-int | LTR |
| NA | ORR1A3-int | LTR |
| 0.8 | ORR1A4 | LTR |
| 0.7 | ORR1B1 | LTR |
| 0.857143 | ORR1E | LTR |
| 0.666667 | RCHARR1 | DNA |
| 1 | RLTR11A2 | LTR |
| NA | RLTR13D3A1 | LTR |
| NA | RLTR13G | LTR |
| NA | RLTR17B_Mm | LTR |
| NA | RLTR19-int | LTR |
| 1 | RLTR1B-int | LTR |
| NA | RLTR20A4 | LTR |
| NA | RLTR20D | LTR |
| NA | RLTR9E | LTR |
| 1 | RMER12C | LTR |
| NA | RMER13B | LTR |
| 1 | RMER16-int | LTR |
| NA | RMER17B2 | LTR |
| 1 | RMER17C | LTR |
| NA | RMER19B | LTR |
| NA | RMER19C | LTR |

| | | |
|---|---|---|
| 0.547619 | RMER1A | Retroposon |
| 0.666667 | RMER1C | Retroposon |
| NA | RMER20B | LTR |
| NA | RSINE1 | SINE |
| NA | RodERV21-int | LTR |
| 1 | URR1A | DNA |

*Supplementary Table 4. P-values of the identified RE in the methylome data comparing ON and CT of F2 generation. NA values are due to low individuals.*

| P-value | RE | General type of RE |
|---|---|---|
| NA | B3 | SINE |
| 0.533333 | B3A | SINE |
| 0.628571 | B4 | SINE |
| 0.125541 | B4A | SINE |
| 0.719695 | IAPEz-int | LTR |
| 0.666667 | IAPLTR1_Mm | LTR |
| 0.914286 | IAPLTR1a_Mm | LTR |
| 1 | IAPLTR2_Mm | LTR |
| 1 | IAPLTR2a | LTR |
| NA | IAPLTR2a2_Mm | LTR |
| 1 | IAPLTR2b | LTR |
| 0.690476 | L1Lx_I | LINE |
| 0.533333 | L1Lx_III | LINE |
| 0.333333 | L1Lx_IV | LINE |
| 0.628571 | L1M2 | LINE |

| | | |
|---|---|---|
| 0.666667 | L1M4 | LINE |
| 0.666667 | L1M5 | LINE |
| 1 | L1MA4A | LINE |
| 0.666667 | L1MA6 | LINE |
| 0.4 | L1MB4 | LINE |
| NA | L1MC1 | LINE |
| 0.666667 | L1MD2 | LINE |
| NA | L1MDa | LINE |
| 1 | L1MEc | LINE |
| 0.412698 | L1MdA_I | LINE |
| 0.754579 | L1MdA_II | LINE |
| 0.413586 | L1MdA_III | LINE |
| 1 | L1MdA_V | LINE |
| 0.547619 | L1MdA_VI | LINE |
| 1 | L1MdA_VII | LINE |
| 0.8 | L1MdF_I | LINE |
| 0.818182 | L1MdF_III | LINE |
| 0.17094 | L1MdF_IV | LINE |
| 1 | L1MdF_V | LINE |
| 0.730159 | L1MdFanc_I | LINE |
| 1 | L1MdFanc_II | LINE |
| 0.2 | L1MdGf_II | LINE |
| **0.018182** | **L1MdMus_I** | **LINE** |
| 1 | L1MdMus_II | LINE |
| 0.666667 | L1MdV_II | LINE |
| 1 | L1_Mur1 | LINE |
| 0.329004 | L1_Mur2 | LINE |

| | | |
|---|---|---|
| 0.4 | L1_Rod | LINE |
| 0.114286 | L2a | LINE |
| 1 | L2b | LINE |
| NA | Lx | LINE |
| 1 | Lx2B2 | LINE |
| NA | Lx3A | LINE |
| 0.666667 | Lx3B | LINE |
| 0.666667 | Lx4A | LINE |
| 0.333333 | Lx5 | LINE |
| 0.527273 | Lx5c | LINE |
| 0.431818 | Lx6 | LINE |
| 0.833333 | Lx7 | LINE |
| 0.787879 | Lx8 | LINE |
| 0.333333 | Lx8b | LINE |
| 0.5 | Lx9 | LINE |
| 0.228571 | MERVL-int | LTR |
| 0.4 | MERVL_2A-int | LTR |
| 1 | MMERVK10C-int | LTR |
| 0.2 | MMERVK9C_I | LTR |
| 1 | MMVL30-int | LTR |
| 0.518482 | MTA_Mm | LTR |
| NA | MTB | LTR |
| 1 | MTC | LTR |
| 0.282828 | MTD | LTR |
| 1 | MTE-int | LTR |
| 0.142857 | MTEa | LTR |
| 1 | MuRRS-int | LTR |

| | | |
|---|---|---|
| 1 | ORR1A0 | LTR |
| 0.2 | ORR1A2 | LTR |
| 0.699301 | ORR1B1-int | LTR |
| NA | ORR1B2 | LTR |
| 0.666667 | ORR1C1 | LTR |
| 0.927273 | ORR1C2 | LTR |
| 0.571429 | ORR1D1 | LTR |
| 0.266667 | ORR1D2 | LTR |
| 0.133333 | ORR1F | LTR |
| NA | ORR1G | LTR |
| NA | RLTR10 | LTR |
| 0.7 | RLTR10-int | LTR |
| 0.628571 | RLTR20C1_MM | LTR |
| 1 | RLTR40 | LTR |
| 0.190476 | RLTR4_MM-int | LTR |
| 0.857143 | RMER15 | LTR |
| 0.7 | RMER15-int | LTR |
| NA | RMER19B2 | LTR |
| 0.699134 | RMER1B | Retroposon |
| 0.666667 | RMER30 | DNA |
| NA | RMER4B | LTR |
| 0.704504 | not_RE | . |
| NA | (GTGTGT)n | Simple_repeat |
| NA | B1F | SINE |
| NA | B1_Mm | SINE |
| 1 | ERV3-16A3_I | LTR |
| NA | ERVB4_2-I_MM | LTR |

| | | |
|---|---|---|
| NA | ERVL-int | LTR |
| 1 | ETnERV-int | LTR |
| 1 | ETnERV2-int | LTR |
| 1 | HERV16-int | LTR |
| 1 | IAP-d-int | LTR |
| 0.666667 | IAPEY2_LTR | LTR |
| 0.536797 | IAPEY3-int | LTR |
| 1 | IAPEY5_I | LTR |
| 1 | IAPEy-int | LTR |
| 0.285714 | IAPLTR3-int | LTR |
| 1 | IAPLTR4_I | LTR |
| 0.2 | ID_B1 | SINE |
| 1 | L1M3 | LINE |
| 1 | L1M3c | LINE |
| NA | L1M4b | LINE |
| NA | L1M4c | LINE |
| 0.7 | L1MA4 | LINE |
| NA | L1MA5 | LINE |
| NA | L1MA9 | LINE |
| 1 | L1MB3 | LINE |
| NA | L1MB5 | LINE |
| NA | L1MB7 | LINE |
| NA | L1MB8 | LINE |
| NA | L1MCa | LINE |
| NA | L1ME1 | LINE |
| NA | L1MEg | LINE |
| 0.8 | L1MdV_I | LINE |

| | | |
|---|---|---|
| 0.666667 | L1MdV_III | LINE |
| 0.114286 | L1_Mur3 | LINE |
| 0.4 | L2 | LINE |
| 0.666667 | L2c | LINE |
| 0.8 | L3 | LINE |
| NA | Lx2 | LINE |
| NA | Lx2A | LINE |
| NA | Lx2B | LINE |
| NA | Lx3_Mus | LINE |
| NA | Lx4B | LINE |
| NA | Lx5b | LINE |
| 1 | MER2 | DNA |
| 1 | MIRb | SINE |
| NA | MIRc | SINE |
| 1 | MLT-int | LTR |
| NA | MLT1A | LTR |
| 1 | MLT1A0 | LTR |
| NA | MLT1B | LTR |
| 1 | MLT1D | LTR |
| NA | MLT1H | LTR |
| 0.666667 | MLT1K | LTR |
| NA | MMERVK10D3_I | LTR |
| 1 | MMERVK9E_I | LTR |
| NA | MT2A | LTR |
| NA | MT2B1 | LTR |
| 1 | MT2B2 | LTR |
| 0.333333 | MTB_Mm | LTR |

| | | |
|---|---|---|
| NA | MTC-int | LTR |
| 0.666667 | MTE2a | LTR |
| 1 | MTE2b | LTR |
| 0.666667 | MTEb | LTR |
| NA | MYSERV-int | LTR |
| 1 | MYSERV16_I | LTR |
| NA | MuRRS4-int | LTR |
| NA | MurERV4-int | LTR |
| NA | ORR1A1-int | LTR |
| 1 | ORR1A3-int | LTR |
| 0.333333 | ORR1A4 | LTR |
| 1 | ORR1B1 | LTR |
| 0.4 | ORR1E | LTR |
| NA | RCHARR1 | DNA |
| NA | RLTR11A2 | LTR |
| 1 | RLTR13D3A1 | LTR |
| 1 | RLTR13G | LTR |
| 1 | RLTR17B_Mm | LTR |
| 0.666667 | RLTR19-int | LTR |
| NA | RLTR1B-int | LTR |
| 1 | RLTR20A4 | LTR |
| NA | RLTR20D | LTR |
| 1 | RLTR9E | LTR |
| NA | RMER12C | LTR |
| 0.666667 | RMER13B | LTR |
| NA | RMER16-int | LTR |
| NA | RMER17B2 | LTR |

| | | |
|---|---|---|
| NA | RMER17C | LTR |
| 1 | RMER19B | LTR |
| 1 | RMER19C | LTR |
| 0.666667 | RMER1A | Retroposon |
| 0.8 | RMER1C | Retroposon |
| NA | RMER20B | LTR |
| 0.5 | RSINE1 | SINE |
| NA | RodERV21-int | LTR |
| NA | URR1A | DNA |
